# Mathematical Modeling of Thyroid Homeostasis: Implications for the Allan-Herndon-Dudley Syndrome

**DOI:** 10.1101/2022.01.24.476744

**Authors:** Tobias M. Wolff, Carina Veil, Johannes W. Dietrich, Matthias A. Müller

**Author notes:** E-mail:{, }. **Corresponding Author:** Tobias M. Wolff.

## Abstract

**Objective:** A mathematical model of the pituitary-thyroid feedback loop is extended to deepen the understanding of the Allan-Herndon-Dudley syndrome (AHDS).

**Background:** The AHDS is characterized by unusual thyroid hormone concentrations and a mutation in the *SLC16A2* gene encoding for the monocarboxylate transporter 8 (MCT8). This mutation leads to a loss of thyroid hormone transport activity. One hypothesis to explain the unusual hormone concentrations of AHDS patients is that due to the loss of thyroid hormone transport activity, thyroxine (*T*_4_) is partially retained in thyroid cells.

**Methods:** This hypothesis is investigated by extending a mathematical model of the pituitary-thyroid feedback loop to include a model of the net effects of membrane transporters such that the thyroid hormone transport activity can be considered. Two modeling approaches of the membrane transporters are employed: on the one hand a nonlinear approach based on the Michaelis-Menten kinetics and on the other hand its linear approximation. The unknown parameters are identified through a constrained parameter optimization.

**Results:** In dynamic simulations, damaged membrane transporters result in a retention of *T*_4_ in thyroid cells and ultimately in the unusual hormone concentrations of AHDS patients. The two different modeling approaches lead to similar results.

**Conclusion:** The results support the hypothesis that a partial retention of *T*_4_ in thyroid cells represents one mechanism responsible for the unusual hormone concentrations of AHDS patients. Moreover, our results suggest that the retention of *T*_4_ in thyroid cells could be the main reason for the unusual hormone concentrations of AHDS patients.

## Introduction

The AHDS is a rare and severe disease which was first described in 1944 by Allan, Herndon and Dudley (Allan et al., 1944). Patients suffer from different symptoms like, e.g., hypotonia, primitive reflexes, scoliosis, muscular hypoplasia and dystonia (Groeneweg et al., 2020).

A first key observation regarding AHDS patients are low free *T*_4_ (*FT*_4_), slightly elevated thyroid stimulating hormone (*TSH*) and high free triiodothyronine (*FT*_3_) concentrations compared to healthy individuals (Groeneweg et al., 2020; Schwartz et al., 2005). A second key observation of AHDS patients are mutations in the *SLC16A2* gene encoding for the monocarboxylate transporter 8 (MCT8) (Dumitrescu et al., 2004; Friesema et al., 2004), which is a specific thyroid hormone transporter (Friesema et al., 2003). A mutation in the related gene often goes along with a complete loss of thyroid hormone transport activity (Friesema et al., 2010).

To elucidate the exact mechanisms that lead to the altered hormone concentrations of this disease, several studies have been made with Mct8 knockout (KO) mice (Trajkovic-Arsic et al., 2010a,b; Dumitrescu et al., 2006). These mice miss the MCT8 and are therefore suitable for investigations related to this disease (Dumitrescu et al., 2006). Furthermore, the hormone concentrations of Mct8 KO mice are strongly similar to those of AHDS patients (Wirth et al., 2009).

Trajkovic-Arsic et al. (2010a) documented that the thyroids of Mct8 KO mice contain approximately a 3-fold elevation of *T*_4_ compared to wild-type littermates. Based on this finding, the hypothesis was made that one mechanism responsible for the unusual hormone concentrations of AHDS patients is a partial retention of *T*_4_ in thyroid cells (Mueller and Heuer, 2012). In this case, more *T*_4_ would be converted into *T*_3_ by thyroidal 5’-deiodinase type I (D1) (Mueller and Heuer, 2012). The result would be that the serum *FT*_4_ concentrations of AHDS patients are lower compared to healthy individuals. Assuming that the *T*_3_ release of the thyroid is not harmed (Trajkovic-Arsic et al., 2010a), the serum *FT*_3_ concentrations of AHDS patients would be higher compared to healthy individuals. The feedback signal of *T*_4_ at the pituitary would induce a higher serum concentration of *TSH*.

If there were no additional mechanisms in the genesis of the unusual hormone concentrations than the one described by Mueller and Heuer (2012), one would expect that athyroid Mct8 KO mice receiving exogenous thyroid hormone supply do not show the unusual hormone concentrations. In athyroid mice no retention of *T*_4_ in the thyroid can take place and thus the effects of this mechanism should not exist. Investigations with athyroid Mct8 KO mice reveal that it is possible to establish normal *T*_3_ concentrations by exogenous *T*_4_ supply (Trajkovic-Arsic et al., 2010a). In turn, the *T*_4_ concentrations remain very low (Trajkovic-Arsic et al., 2010a). This observation indicates that the retention of *T*_4_ in the thyroid gland does contribute to the high *T*_3_ concentrations but not to the low *T*_4_ concentration (Mueller and Heuer, 2012). Therefore, Mueller and Heuer (2012) draw the conclusion that additional mechanisms must contribute to the unusual hormone concentrations. Particularly, they suggest a renal contribution, since *T*_4_ accumulates in the kidney and the activity of D1 inside the kidney is increased for Mct8 KO mice (Trajkovic-Arsic et al., 2010b).

In this work, we investigate the mechanisms leading to the unusual hormone concentrations of AHDS patients by means of a mathematical model of the pituitary–thyroid feedback loop and dynamic simulations. In detail, we further extend the model originally developed by Dietrich (2001); Dietrich et al. (2004); Berberich et al. (2018) to include membrane transporters between the thyroid gland and the periphery. Our results indicate that damaged membrane transporters, i.e., a loss of thyroid hormone transport activity leads to an increased *T*_4_ content in thyroid cells and ultimately to the unusual hormone concentrations that are measured at AHDS patients. These results lead to a partially different conclusion compared to the suggestions of Mueller and Heuer (2012), since they suggest that the entirety of the unusual hormone concentrations of AHDS patients be explained by a retention of *T*_4_ in thyroid cells, without additional renal contribution.

## Methods

First, we introduce the mathematical model of the pituitary-thyroid feedback loop and its extension to membrane transporters. Second, we propose a method to identify the unknown parameters, namely the constrained parameter optimization.

### Extended Model

In general, a mathematical model describes a system by taking into account the available knowledge about the underlying cause-effect relationships. To understand the fundamental principles of the model used throughout this work, a simplified block diagram of it is illustrated in Figure 1, whereas a more detailed description is given in Figure 2. The basic principle is that *TSH* stimulates the production of *T*_3_ and *T*_4_ at the thyroid gland. By means of D1 and 5’deiodinase type II (D2), *T*_4_ is converted into *T*_3_ in peripheral organs like the liver and the kidney. The production of *TSH* at the pituitary is inhibited by *T*_4_, whereas the production of *TSH* is stimulated by thyrotropin-releasing hormone (*TRH*). The detailed mathematical model used in this work describes the cause-effect relationships by six nonlinear differential equations visible in Figure 2. A detailed state of the art description of the mathematical model as developed by Dietrich (2001); Dietrich et al. (2004); Berberich et al. (2018) is given in Section S1 of the Supplementary Material, so that this paper is as self-contained as possible. In the main part, we will emphasize on the extension of the model, conducted within this work.

**Figure 1:**
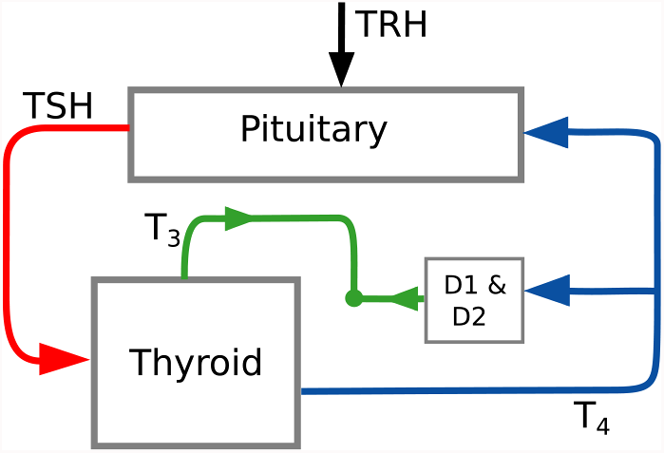
Simplified block diagram of the pituitary-thyroid feedback loop. This diagram illustrates the main structure of the applied mathematical model. The detailed model is illustrated in Figure 2. The parameter D2 and the variable *TRH* denote the 5’ deiodinase type II and the thyrotropin-releasing hormone, respectively.

**Figure 2:**
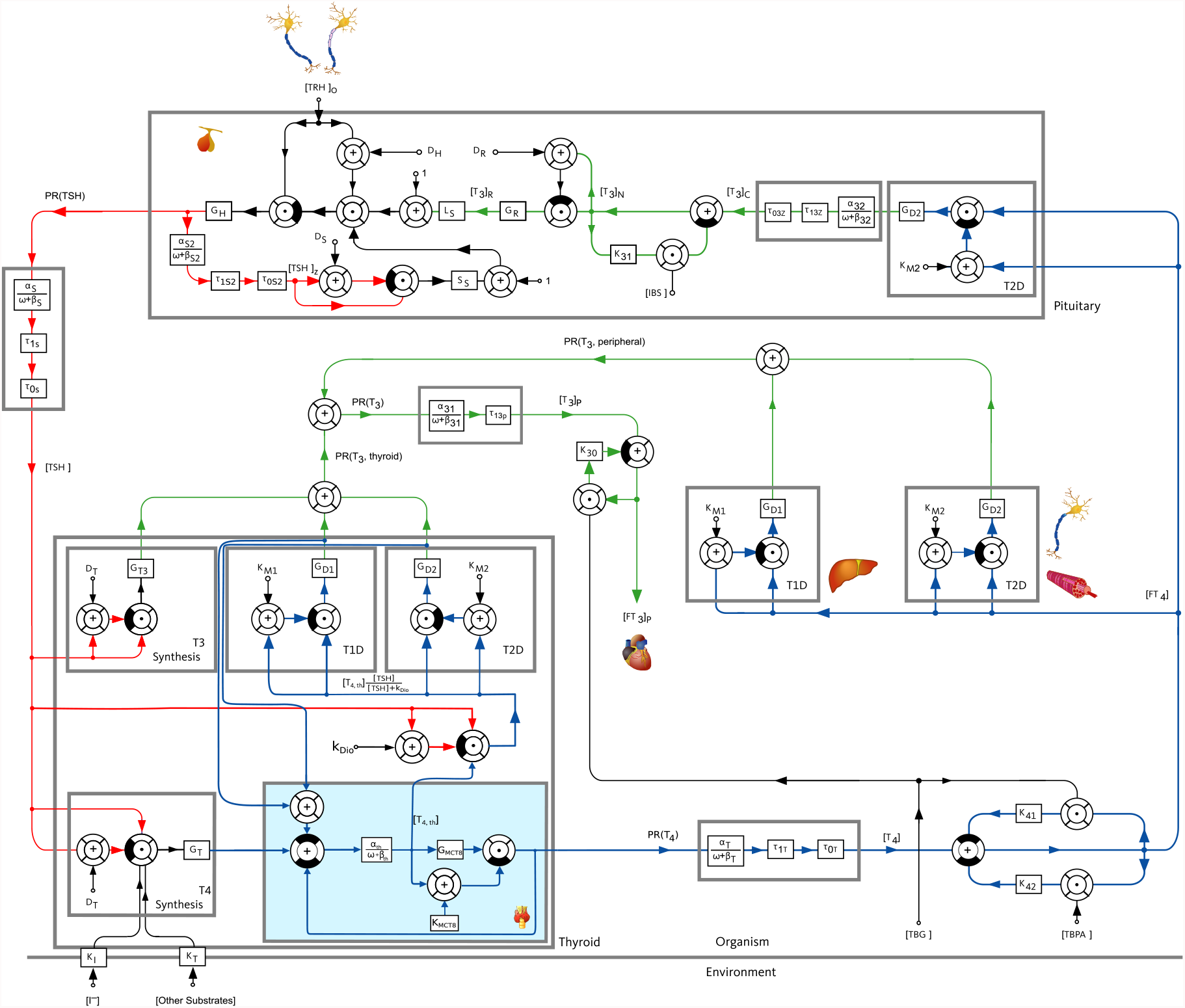
Block diagram of the pituitary-thyroid feedback loop including membrane transporters, extended from Dietrich (2001); Dietrich et al. (2004); Berberich et al. (2018). The extension presented in this paper is shown with a slight blue background color in the block “Thyroid”. The numerical parameter values are mostly taken from Dietrich (2001); Berberich et al. (2018). The values of *G*_*D*1_, *G*_*T* 3_ and *G*_*MCT* 8_ are identified through a constrained parameter optimization using real hormone measurements from Dietrich (2001).

The investigations of the effects of damaged membrane transporters on the hormone concentrations necessitate a representation of the net effects of the membrane transporters in the mathematical model of the pituitary-thyroid feedback loop from Dietrich (2001); Dietrich et al. (2004); Berberich et al. (2018). For simplicity, we will call the modeled net effects of the membrane transporters “Michaelis-Menten modeling” (or “linear modeling”) of the membrane transporters, even though we do not refer to a specific concentration or state variable but to the net effects.

We incorporate membrane transporters between the thyroid gland and the periphery for *T*_4_ (illustrated in Figure 2 by a slight blue background color) in order to analyze the mentioned hypothesis. Obviously, one could incorporate membrane transporters at a number of different locations in the model, but this would further complicate the model and result in difficulties in the identification of the additional parameters of the membrane transporters.

Moreover, further incorporations of the effects of MCT8 deficiency are not crucial to pursue the objective of analyzing the hypothesis of Mueller and Heuer (2012) that a retention of *T*_4_ in thyroid cells represents one mechanism responsible for the unusual hormone concentrations of AHDS patients.

Following the common modeling approach of enzyme/substrate reactions based on the well known Michaelis-Menten kinetics (Murray, 2007), the functionality of the membrane transporters is considered by

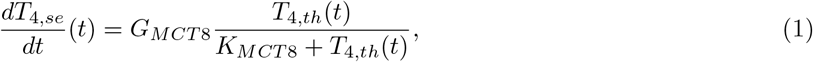

where *T*_4,*th*_ represents the *T*_4_ concentration in thyroid cells and *T*_4,*se*_ the *T*_4_ in the blood circulation. *G*_*MCT* 8_ stands for the maximal activity of the MCT8 and *K*_*MCT* 8_ for the Michaelis-Menten constant of the transport process. Our model reflects the organ level. Hence, the modeled membrane transporters can be interpreted as the cumulated effect of single membrane transporters on cell level.

The incorporation of the membrane transporters into the already existing model (Berberich et al., 2018) is done by an introduction of a new state, named *T*_4,*th*_. Its differential equation is defined by

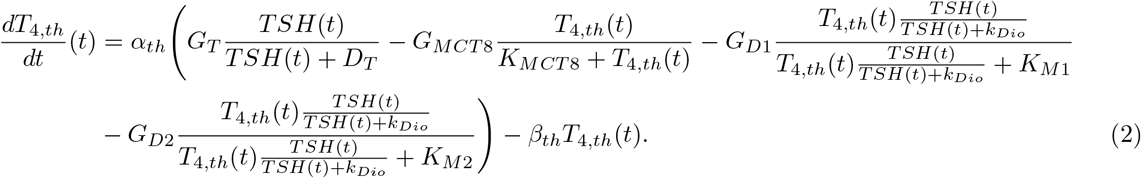

The first term describes the production rate of *T*_4_, with *G*_*T*_ being the secretory capacity of the thyroid gland and *D*_*T*_ representing the damping constant at the thyroid gland (compare Section S1 of the Supplementary Material for a more detailed explanation of the model and the meaning of the parameters). The second term stands for the part of *T*_4_, which is transported out of the thyroid cells. The remaining terms represent the thyroidal conversion of *T*_4_ into *T*_3_ by D1 and D2, where the maximal activity of D1/D2 is denoted by *G*_*D*1_*/G*_*D*2_ and the dissociation constant of D1/D2 by *K*_*M*1_/*K*_*M*2_, respectively (compare Berberich et al. (2018) for a detailed discussion of the *TSH*-*T*_3_ shunt).

This conversion rate is considered positively in the differential equation of *T*_3*p*_, equation (S3) of the Supplementary Material. Thus, this conversion rate must be considered negatively in the differential equation of *T*_4,*th*_^1^. The constants *α*_*th*_ and *β*_*th*_ are the dilution factor and the clearance exponent for *T*_4,*th*_. Furthermore, the differential equation of *T*_4_ changes to

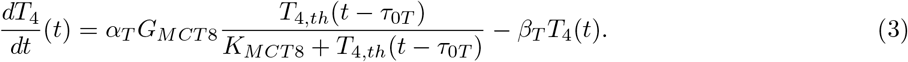

The parameters *α*_*T*_ and *β*_*T*_ are again the dilution factor and the clearance rate constant, respectively, for *T*_4_. The dead time *τ*_0*T*_ is introduced in order to account for diffusion processes. Compared to Berberich et al. (2018), the production rate of the peripheral *T*_4_ does no longer correspond to *G*_*T*_ *TSH/*(*D*_*T*_ + *TSH*), but to the part of *T*_4,*th*_ which is transported out of the thyroid cells.

The numerical values of the majority of the model’s parameters can be taken from Dietrich (2001); Dietrich et al. (2004); Berberich et al. (2018). They have been determined experimentally or derived from known quantities like the half-life of the hormone concentrations. However, the parameters *G*_*D*1_ and *G*_*T* 3_^2^ must be re-identified by fitting them to real measurements of hormone concentrations, because the previous versions of the model did not consider membrane transporters (Dietrich, 2001; Dietrich et al., 2004; Berberich et al., 2018).

In addition, the introduced maximal activity of the MCT8 (*G*_*MCT* 8_) must also be fitted to real measurements. In turn, the introduced parameter *K*_*MCT* 8_ was determined experimentally by Friesema et al. (2003) and its value is applied here. This is a meaningful approach, since the Michaelis-Menten constant remains the same for a specific transporter/substrate process (Berg et al., 2002). To find the numerical parameter values, we neglect the age dependence of the *T*_3_ concentrations of AHDS patients, as documented in Groeneweg et al. (2020) and the age dependence of the *T*_4_ content in thyroid cells, documented in Di Cosmo et al. (2010). Even though such a consideration would certainly be advantageous, it is not indispensable and would further add complexity to the model.

The factor *α*_*th*_ is defined as the inverse of the volume of distribution of *T*_4_ in the thyroid gland. We choose 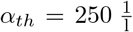, which corresponds to a volume of distribution of 4 ml. This volume of distribution is based on the assumption that the intracellular parts of the thyroid gland make up one third of the whole volume of the thyroid gland. Furthermore, we choose the clearance exponent for *T*_4_ in the thyroid gland as *β*_*th*_ = 4.4 *·* 10^−6^ s^−1^ corresponding to a half-life of *T*_4_ in the thyroid gland of 44 h, a value determined in Di Cosmo et al. (2010) for rat thyroids. We want to emphasize that the exact numerical values of *α*_*th*_ and *β*_*th*_ do not considerably influence our results. Even if the true values differ to some extent from the ones that we suggest here, our main results (see next section) remain the same, i.e., that the hormome levels of AHDS patients can be explained by damaged membrane transporters. A summary of the entire numerical parameter values can be found in Section S6 in the Supplementary Material.

### Parameter Identification

The parameter identification of *G*_*D*1_, *G*_*T* 3_ and *G*_*MCT* 8_ is done through a constrained parameter optimization. The idea is to find the optimal configuration of parameters with respect to a cost function by adhering to the system dynamics. We chose the cost function as the normalized quadratic error between real measured hormone concentrations and the steady-state hormone concentrations computed by the model. The adherence to the system dynamics is realized by forcing the solution of the optimization problem to fulfill the steady-state equations. One encountered challenge regarding the parameter identification were the differences in the magnitudes of the steady-state equations, e.g., the value of *FT*_3_ is of order 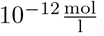, whereas the order of *TSH* is 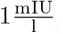.

This can pose a problem, since we want that the steady-state equations are fulfilled by the solution of the optimization problem up to some tolerance value. Consider examplarily that the solution of the optimization problem fulfills all steady-state equations up to the order of magnitude 10^−12^, which would be a satisfactory result regarding *TSH*, but an unsatisfactory result regarding *FT*_3_, since an error of 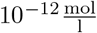 makes a substantial difference.

To overcome this problem, a state transformation was performed which renders all steady-state equations to the same order of magnitude. The exact procedure of the identification was as follows: first, we identified the parameters *G*_*D*1_, *G*_*T* 3_ and *G*_*MCT* 8_ for healthy individuals. To this end, we used the real hormone measurement data of 81 untreated healthy individuals from Dietrich (2001). Second, the identified parameters of *G*_*D*1_ and *G*_*T* 3_ are held constant and only the value of *G*_*MCT* 8_ is fitted to real measurement hormone data for AHDS patients. In this case, we used the documented mean values of *TSH* and *FT*_4_ of Groeneweg et al. (2020) and the mean *FT*_3_ value of Schwartz et al. (2005)^3^. The exact mathematical formulation of the parameter identification is shown in Section S2 in the Supplementary Material. Since the mentioned parameters are identified using real hormone concentration measurements from different subjects, the models for healthy individuals and for AHDS patients can be interpreted as models of a generic euthyroid subject and a generic AHDS patient, respectively. After identifying the parameters with the described procedure, one can perform dynamic simulations in order to evaluate the hormone concentrations of healthy individuals and AHDS patients. We chose (arbitrarily) a simulation length of 30 days in order to illustrate the course of the hormone concentrations for a period of one month.

When taking a closer look on the expression of (1), one recognizes that a linear approximation of the term is possible if *K*_*MCT* 8_ *>> T*_4,*th*_. By defining the constant *k*_*l*_ = *G*_*MCT* 8_*/K*_*MCT* 8_, equation (1) becomes

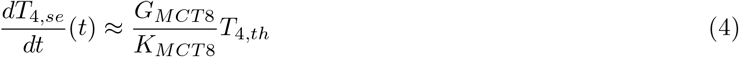

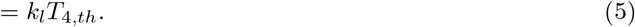

This means that we can model the functionality of the membrane transporters linearly if the mentioned condition is fulfilled. In this case, equations (2) and (3) must be changed accordingly. The procedure of the parameter identification remains the same. The only difference is that we have to identify *k*_*l*_ instead of *G*_*MCT* 8_.

As an additional contribution, we performed a local stability analysis. So far, this was not done for any other mathematical model of the pituitary-thyroid feedback loop (Dietrich, 2001; Dietrich et al., 2004; Berberich et al., 2018). A stability analysis is interesting for the highly perturbed pituitary-thyroid feedback loop system of AHDS patients. It helps to answer the question whether the feedback loop is stable for damaged membrane transporters. The presentation of the employed method and of the corresponding results are given in the Supplementary Material Section S4. This analysis reveals asymptotic stability of the equilibrium hormone concentrations.

## Results

First, the results of the parameter identification and of the dynamic simulations for the Michaelis-Menten modeling of the membrane transporters are presented. Second, the analogous results for the linear modeling of the membrane transporters are shown.

### Michaelis-Menten Modeling

The results of the parameter identification are shown in Table 1. Regarding healthy individuals, the initial guess of *G*_*D*1_ corresponds to the one determined by Berberich et al. (2018). Since the value of *G*_*MCT* 8_ was never measured nor identified previously, its initial guess is set equal to *K*_*MCT* 8_. The initial guesses of the hormone concentrations correspond to the mean values of the real measured hormone concentrations from Dietrich (2001) (*T*_4,*th*_ corresponding to the formula of Berberich et al. (2018)). For the optimization problems we tested different initial guesses, since we solve a non-convex optimization problem with a local solver (for details see the Supplementary Material Section S3). Concerning healthy individuals, different initial guesses influence the optimal configuration of the parameters. In detail, different final values of *G*_*D*1_, *G*_*T* 3_ and *G*_*MCT* 8_ lead to the same optimal values of *FT*_3_, *FT*_4_ and *TSH* values, thus, to the same value of the cost function. This means that the problem is not uniquely identifiable. A detailed analysis of this observation is presented in Section S3 in the Supplementary Material. Here, we chose a configuration of parameters that leads to the relationship that approximately 20% of the *T*_3_ is produced inside the thyroid and approximately 80% in the periphery, as proposed by Pilo et al. (1990); Bianco et al. (2002). This configuration is reached with an initial guess of 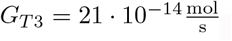.

**Table 1:**
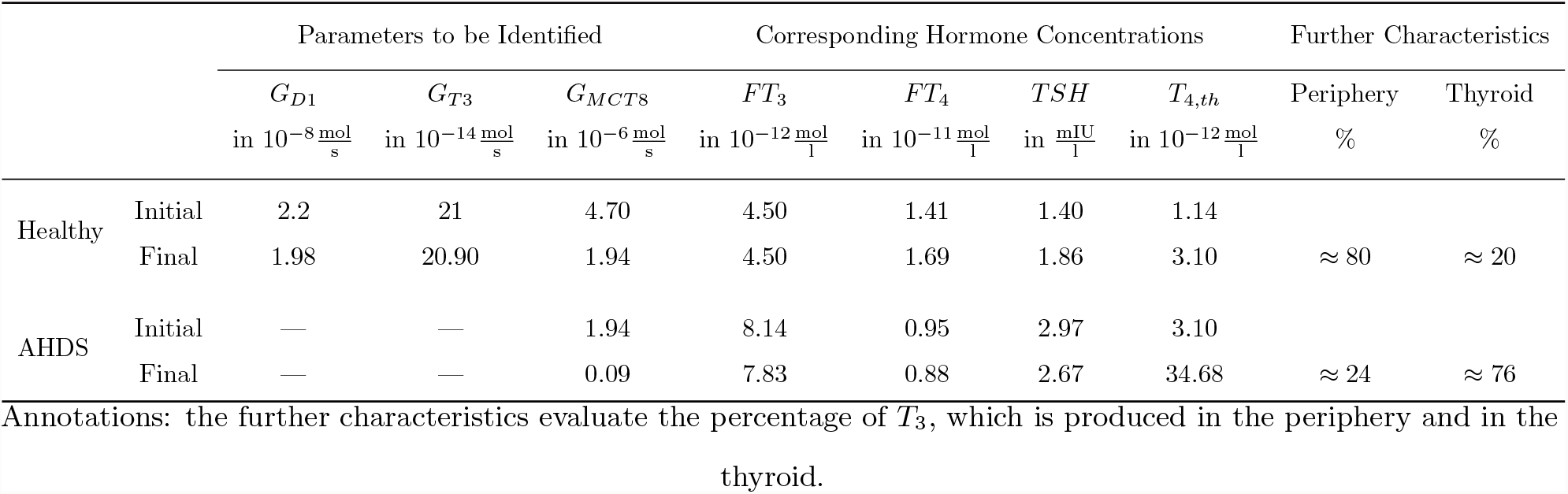
Results of the parameter identification for the Michaelis-Menten modeling of membrane transporters with the constrained parameter optimization

|  |  | Parameters to be Identified |  |  | Corresponding Hormone Concentrations |  |  |  | Further Characteristics |  |
| --- | --- | --- | --- | --- | --- | --- | --- | --- | --- | --- |
| | | $G_{D1}$<br>in $10^{-8} \frac{\text{mol}}{\text{s}}$ | $G_{T3}$<br>in $10^{-14} \frac{\text{mol}}{\text{s}}$ | $G_{MCT8}$<br>in $10^{-6} \frac{\text{mol}}{\text{s}}$ | $FT_3$<br>in $10^{-12} \frac{\text{mol}}{\text{l}}$ | $FT_4$<br>in $10^{-11} \frac{\text{mol}}{\text{l}}$ | $TSH$<br>in $\frac{\text{mIU}}{\text{l}}$ | $T_{4,th}$<br>in $10^{-12} \frac{\text{mol}}{\text{l}}$ | Periphery<br>% | Thyroid<br>% |
| Healthy | Initial | 2.2 | 21 | 4.70 | 4.50 | 1.41 | 1.40 | 1.14 |  |  |
| | Final | 1.98 | 20.90 | 1.94 | 4.50 | 1.69 | 1.86 | 3.10 | $\approx 80$ | $\approx 20$ |
| AHDS | Initial | — | — | 1.94 | 8.14 | 0.95 | 2.97 | 3.10 |  |  |
| | Final | — | — | 0.09 | 7.83 | 0.88 | 2.67 | 34.68 | $\approx 24$ | $\approx 76$ |

Annotations: the further characteristics evaluate the percentage of $T_3$ , which is produced in the periphery and in the thyroid.

Regarding AHDS patients, the initial guesses of *FT*_3_, *FT*_4_ and *TSH* correspond to the mean measured hormone concentrations (Groeneweg et al., 2020; Schwartz et al., 2005). The initial guesses of *G*_*MCT* 8_ and *T*_4,*th*_ are equal to the identified values of healthy individuals. For AHDS patients, the parameter *G*_*MCT* 8_ can be identified uniquely. Different initial guesses do not influence the final result. This difference to the identification of the parameters for healthy individuals is due to the fact that we only identify one unknown parameter, namely *G*_*MCT* 8_, and not three (*G*_*D*1_, *G*_*T* 3_ and *G*_*MCT* 8_) as it is the case for healthy individuals. Therefore, we do not need any additional information to identify the value of *G*_*MCT* 8_ for AHDS patients.

Once the unknown parameter values are determined, dynamic simulations can be performed. The course of the hormone concentrations of *TSH, FT*_4_, *FT*_3_ and *T*_4,*th*_ are given in Figure 3, where a sinusoidal *TRH* input was used. Regarding the concentration of *TSH*, one can see that the hormone concentrations of AHDS patients are slightly higher compared to the ones of healthy individuals. The course of *FT*_3_ reveals a far higher concentration for AHDS patients than for healthy individuals. Concerning the concentration of *FT*_4_, AHDS patients have a lower concentration than healthy individuals. As mentioned previously, the introduced state *T*_4,*th*_ describes the *T*_4_ content in thyroid cells. Figure 3 shows that the *T*_4_ content in thyroid cells is far higher (approximately an 11-fold increase) for AHDS patients than for healthy individuals.

**Figure 3:**
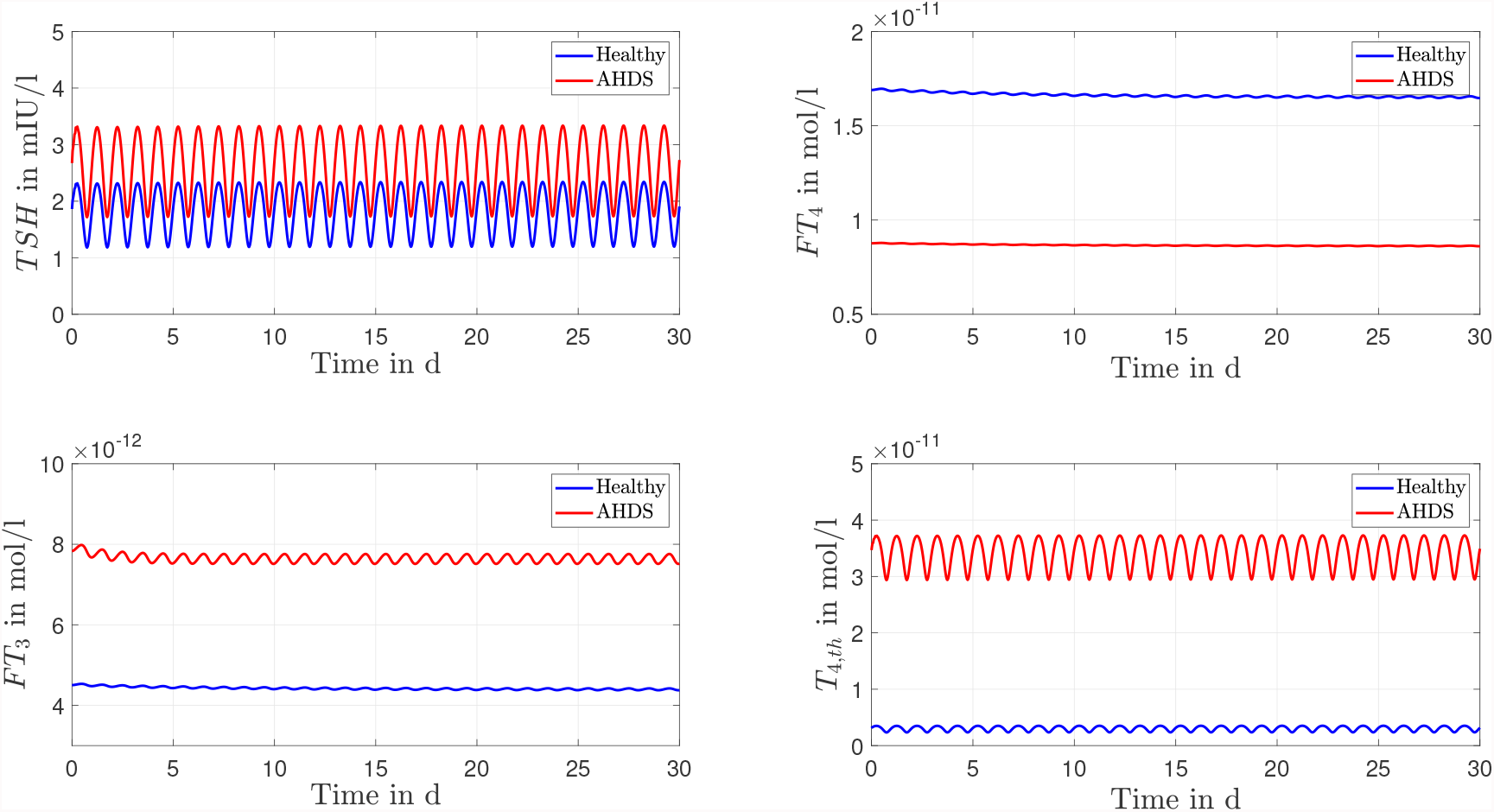
Results of the dynamic simulations for the Michaelis-Menten modeling of the membrane transporters. Most of the numerical parameter values are based on the suggestions of Dietrich (2001); Berberich et al. (2018). The remaining unknown parameters of the model are identified through a constrained parameter optimization and shown in Table 1.

### Linear Modeling

We now focus on the results of the linear approximation of the membrane transporters. As can be seen from Table 1, the assumption *K*_*MCT* 8_ = 4.7 *·* 10^−6^ *>> T*_4,*th*_ = 3.10 *·* 10^−12^ is fulfilled. Therefore, the same analysis as in the previous subsection is performed with the linear approximation of the membrane transporters. The results of the parameter identification are shown in Table 2. Again, the parameters cannot be identified uniquely for healthy individuals: different final values of *G*_*D*1_, *G*_*T* 3_ and *k*_*l*_ lead to the same final values of *TSH, FT*_4_ and *FT*_3_. The final values of the *FT*_3_, *FT*_4_ and *TSH* concentrations do not differ from the Michaelis-Menten modeling approach. The final value of *k*_*l*_ corresponds approximately to *G*_*MCT* 8_/*K*_*MCT* 8_ (compare Table 1) for healthy individuals. Regarding AHDS patients, we can identify the unknown parameter *k*_*l*_ uniquely. Different initial guesses do not influence the final result. The explanation is analogous to the Michaelis-Menten modeling of the membrane transporters.

**Table 2:**
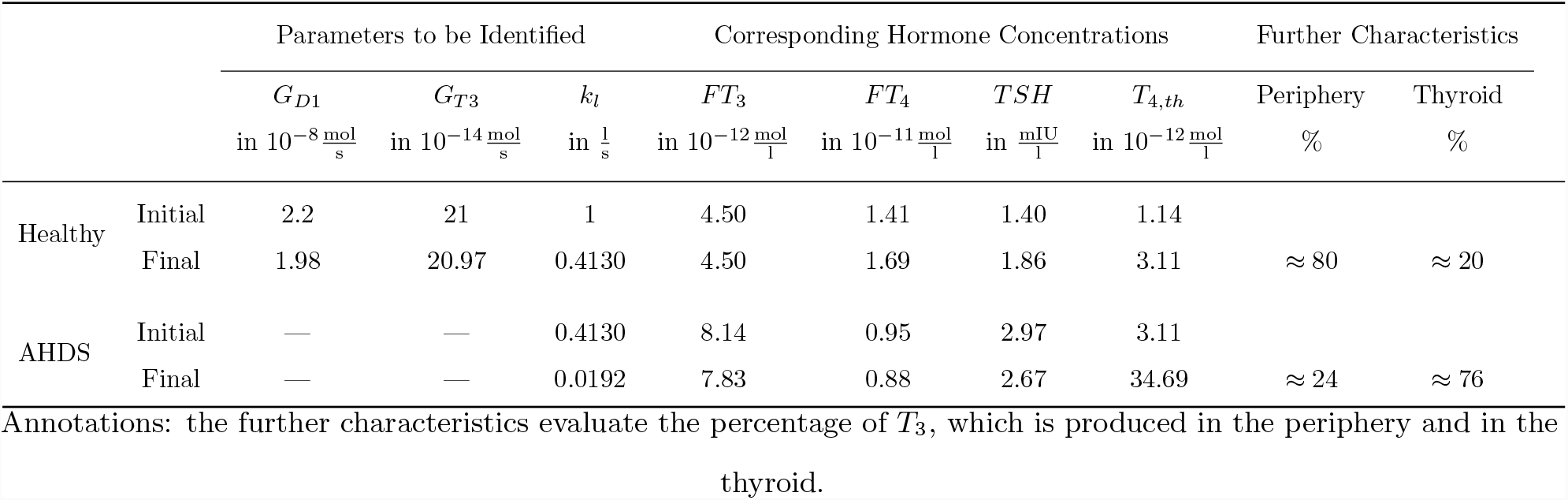
Results of the parameter identification for the linear modeling of the membrane transporters with the constrained parameter optimization approach

Again, one can perform dynamic simulations with the model considering the linearly approximated membrane transporters. The courses of the hormone concentrations are given in Figure 4. One can see that there is virtually no difference in the dynamic course of the hormone concentrations between the Michaelis-Menten modeling of the membrane transporters and its linear approximation. Again, the simulated concentrations of *TSH* and *FT*_3_ are higher for AHDS patients compared to healthy individuals. The concentration of *FT*_4_ is lower for AHDS patients than for healthy individuals. The *T*_4_ content in thyroid cells is approximately 11-times higher for AHDS patients compared to healthy individuals.

**Figure 4:**
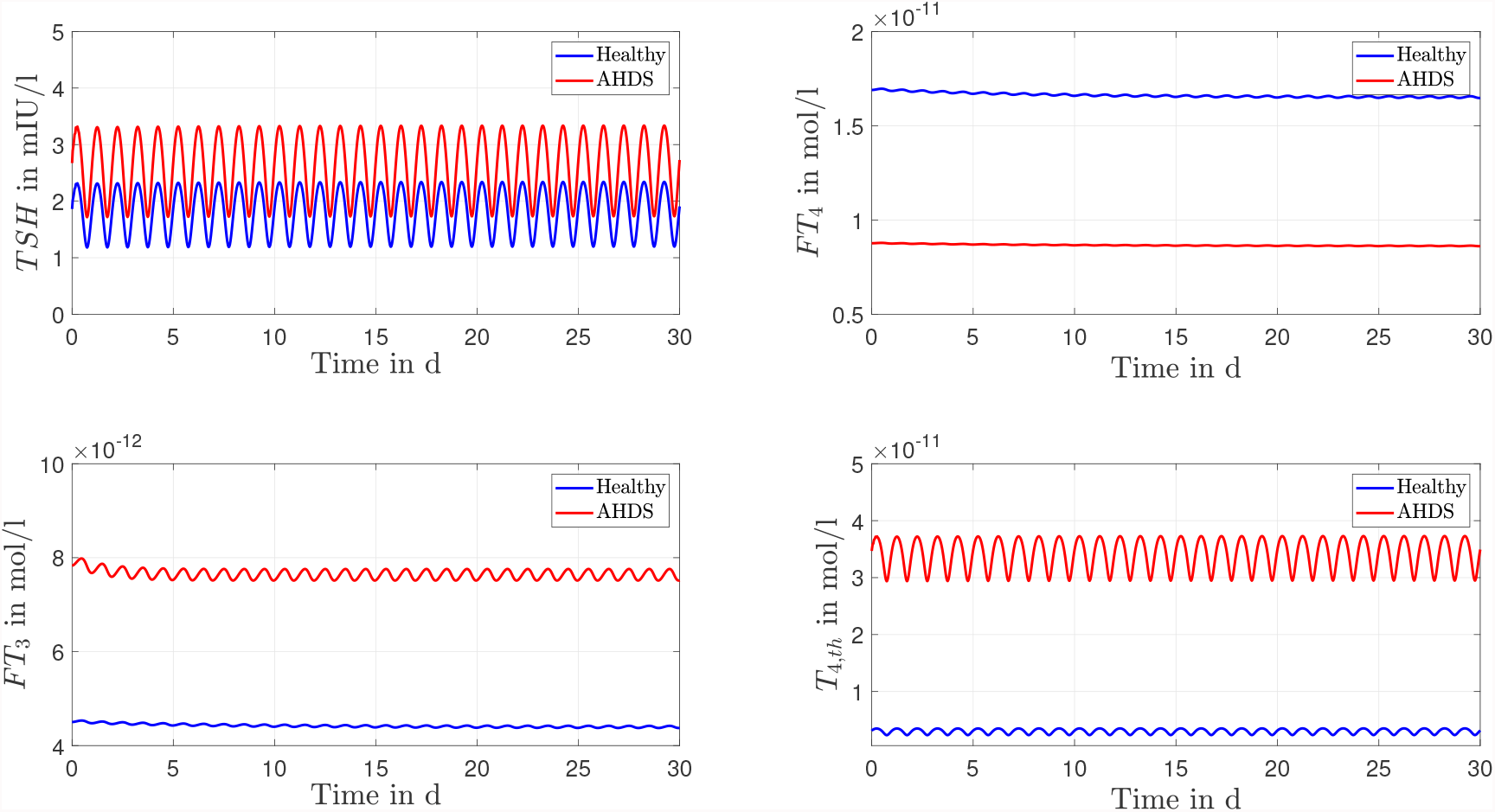
Results of the dynamic simulations of the linear modeling of the membrane transporter. Again, most of the numerical parameter values are based on Dietrich (2001); Berberich et al. (2018). The remaining parameters are identified through a constrained parameter optimization approach and shown in Table 2.

## Discussion

In this section, we discuss the obtained results and place them in a larger context. First, we focus on our main contribution, the investigations around the mechanisms of the AHDS. Second, we analyze the modeling approaches of the membrane transporters and discuss the parameter identification.

### Mechanisms of the AHDS

The documented results of the dynamic simulations (compare Figures 3, 4) clearly demonstrate that the unusual hormone concentrations of AHDS patients are observed in the simulations of the mathematical model when damaged membrane transporters are considered. This result is independent of the modeling approach of the membrane transporters. The simulated hormone concentrations of AHDS patients are additionally in line with the mean measured hormone concentrations presented by Groeneweg et al. (2020) (*TSH* = 2.97 mIU/l, *FT*_4_ = 9.48 *·* 10^−12^mol/l) and Schwartz et al. (2005) (*FT*_3_ = 8.14 *·* 10^−12^mol/l).

One must also keep in mind that the hormone concentrations of healthy individuals should not change substantially, when incorporating membrane transporters. They should remain in the reference range of healthy individuals (*TSH*: 0.5 − 4.5 mIU/l, *FT*_4_: 1.2 − 2.7 *·* 10^−11^ mol/l and *FT*_3_: 3.5 − 6.3 *·* 10^−12^ mol/l (Visser et al. (2013))). The dynamic simulations of the mathematical model reveal (see Figures 3, 4) that the hormone concentrations of healthy individuals remain within the reference range of healthy individuals. Thus, the introduction of the membrane transporters does not impact the usability of the mathematical model for healthy individuals. The incorporation of the membrane transporters extends the possible applications of the mathematical model. Since the concentration of *T*_4,*th*_ does not correspond to a hormone concentration which can be measured with today’s assay technology, it is difficult to evaluate its accordance with human data. Alternatively, the concentration can be compared to studies that are made with mice, e.g., the study of Trajkovic-Arsic et al. (2010a). In this study, an investigation is done regarding the thyroidal *T*_4_ content of Mct8 KO mice. It is reported that the thyroids of these mice contain approximately a 3-fold elevation of *T*_4_ compared to wild-type littermates (Trajkovic-Arsic et al., 2010a). In the presented mathematical model one can interpret the state *T*_4,*th*_ on a high level as *T*_4_ content in thyroid cells. For both modeling approaches of the membrane transporters, the *T*_4_ content in thyroid cells (described by the state *T*_4,*th*_) is approximately 11-times higher for AHDS patients compared to healthy individuals.

Therefore, the mathematical model is in line with the documented observation that the *T*_4_ content in thyroid cells is increased in MCT8 deficiency (Trajkovic-Arsic et al., 2010a). This is an indication that the *T*_4_ content in thyroid cells is not only increased for Mct8 KO mice, but also for AHDS patients. Compared to the results of Trajkovic-Arsic et al. (2010a), one must keep in mind that the difference between the *T*_4_ content in thyroid cells of AHDS patients to healthy individuals is higher in the mathematical model (11-fold increase), than the documented difference of Mct8 KO mice to wild-type littermates (3-fold increase).

At that point we can additionally evaluate the hypothesis that the *T*_4_ retention in thyroid cells represents one important mechanism to the unusual hormone concentrations of AHDS patients (Mueller and Heuer, 2012). In the simulations of this work, damaged membrane transporters go along with an increased *T*_4_ content in thyroid cells and ultimately the unusual hormone concentrations of AHDS patients. Thus, additional evidence is given to the hypothesis of Mueller and Heuer (2012) by means of the mathematical model.

If we take a more precise look on the considerations of Mueller and Heuer (2012), one remarks a small but important difference between their considerations and our results. Mueller and Heuer (2012) assign the high concentration of *FT*_3_ to the retention of *T*_4_ inside the thyroid. In turn, the assumption is made that the low concentrations of *FT*_4_ are due to a renal contribution, as the renal *T*_4_ content is increased in MCT8 deficiency (Mueller and Heuer, 2012; Trajkovic-Arsic et al., 2010b).

The difference to our results is that the simulations lead to the entirety of the unusual hormone concentrations of AHDS patients, including a lower concentration of *FT*_4_. This is an indication that the *T*_4_ retention in thyroid cells does not only represent one mechanism leading to the unusual hormone concentrations of AHDS patients, but could even be fully responsible for these unusual concentrations. In other words, the simulations of the mathematical model suggest that an additional renal contribution might not be necessary to replicate the entirety of the unusual hormone concentrations of AHDS patients.

To evaluate the role of the kidney in the genesis of the unusual hormone concentrations of AHDS patients by means of the mathematical model, an explicit representation of it is necessary. This is currently not the case, because the model merges the effects of the different peripheral organs like the kidney and the liver under one general component, the periphery. Once an explicit representation of the kidney is incorporated into the model, membrane transporters could be considered at the edge of the kidney to the bloodstream, which would allow a more precise investigation of the assumptions from Mueller and Heuer (2012). However, the further refinement of the model usually goes along with more parameters that must be identified. Already in this work it was not possible to identify the parameters uniquely, thus, it will become more and more difficult to find a meaningful configuration of parameters, at least without additional dynamic hormone measurements. Nevertheless, the explicit consideration of the kidney in the mathematical model is an interesting issue for further research.

Finally, we discuss our results with respect to the single case report of Wirth et al. (2009) in which one AHDS patient was examined before and after thyroidectomy. Before thyroidectomy the patient received 75 μg (or 10 μg/kg per day) of levothyroxine (*L*-*T*_4_) to normalize the *TSH* concentration. The exact hormone concentrations were *FT*_4_ ≈ 1.06 *·* 10^−11^ mol/l, *T*_3_ ≈ 7.92 *·* 10^−9^ mol/l and *TSH* = 0.1 mIU/l (compare Figure 2 in Wirth et al. (2009)). After thyroidectomy and 125 μg (or 6 μg/kg per day) of *L*-*T*_4_, the patient’s hormone concentrations were *FT*_4_ ≈ 1.05 *·* 10^−11^ mol/l, *T*_3_ ≈ 4.21 *·* 10^−9^ mol/l and *TSH* = 0.48 mIU/l (again, compare Figure 2 in Wirth et al. (2009)).

Note that the normalization of the *TSH* concentration goes along with a substantially higher concentration of *T*_3_ before thyroidectomy compared to the concentration of *T*_3_ after thyroidectomy. This indicates that a retention of *T*_4_ in thyroid cells could be responsible for the high *T*_3_ concentrations, which is a conclusion in line with the simulation results. The *FT*_4_ concentrations remain approximately constant in both cases. As already mentioned, this points to extrathyroidal mechanisms explaining the low *FT*_4_ concentrations of AHDS patients. In contrast, the simulation result suggest that such extrathyroidal events do not have to be present in order to replicate the hormone concentrations, which seems to be a contradiction at first sight. However, since we do not consider the kidney explicitly in the mathematical model, we also do not exclude extrathyroidal mechanisms from possibly contributing to the unusual hormone concentrations. This aspect can currently not be answered by means of the model, since the kidney is not considered explicitly.

Furthermore, future work could focus on other phenomena related to MCT8 deficiency that were not treated in the context of this work. For example, in Mct8/D1 double KO mice, a partial normalization of thyroid hormone concentrations takes place (compare Liao et al. (2011)). An analysis whether the same observation can be made exploiting the mathematical model of the pituitary-thyroid feedback loop would potentially further deepen the knowledge regarding the AHDS.

### Modeling approaches

Two possibilities to model the membrane transporters are presented in this paper. The Michaelis-Menten modeling of the membrane transporters follows a common approach to model transporter/substrate reactions (Alberts, 2015). A saturation of the transported *T*_4_ can take place, when no MCT8 is available, i.e., *T*_4,*th*_ *>> K*_*MCT* 8_ in equation (1). The linear modeling of the membrane transporters has an appealing simplicity. The intuitive idea that a certain percentage (*k*_*l*_) of *T*_4_ is transported out of thyroid cells is easy to understand. However, this modeling approach is only applicable for a specific range of *T*_4,*th*_, namely as long as *T*_4,*th*_ *<< K*_*MCT* 8_. If this is not the case, the linear approximation is not valid anymore.

In our case, the necessary requirement in order to model the functionality of the membrane transporters linearly is fulfilled (*K*_*MCT* 8_ = 4.7 *·* 10^−6^ >> 3.10 *·* 10^−12^ = *T*_4,*th*_). The results of the parameter identification demonstrate that this approximation is meaningful. This can be seen in the following way. For healthy individuals the term *G*_*MCT* 8_*/K*_*MCT* 8_ ≈ 0.41 of the Michaelis-Menten modeling is approximately equal to *k*_*l*_ ≈ 0.41 of the linear modeling of the membrane transporters (compare final values in Tables 1 and 2). The same holds true for the final values of AHDS patients: *G*_*MCT* 8_*/K*_*MCT* 8_ ≈ 0.02, *k*_*l*_ ≈ 0.02. Given these results, there might be no substantial advantage applying the Michaelis-Menten modeling of the membrane transporters over the linear modeling of the membrane transporters. In conclusion, the application of the linear modeling of the membrane transporters reduces the complexity of the model and leads to similar results.

The applied mathematical model exploits parameters that were determined for humans as well as parameters that were determined for rodents, even though some aspects of the thyroid homeostasis are different. The model does not aim for an exact representation of humans which would be impossible due to an inter-variability of the parameters even for humans only. However, note that it is possible to obtain an understanding of the cause-effect relationships even for these “generic” parameter values (that do not correspond to one individual human subject). Namely, the qualitative behavior that is obtained from simulating the model is the same for different parameter values. Mathematically, this can be shown by a sensitivity analysis for the parameters, compare Berberich et al. (2018). So even if the parameters do not all correspond to the true parameters of a (one individual) human subject, the observed phenomena are still representative for the cause-effect relationships in the human HPT axis.

So far, the loss of thyroid hormone transport activity is only considered at the thyroid gland for *T*_4_. Investigating how MCT8 deficiency affects the complete pituitary-thyroid feedback loop (i.e., considering a loss of thyroid hormone transport activity at further locations) is an interesting topic for future research.

### Parameter Identification

In the context of the parameter identification, we considered the identified values of *G*_*D*1_ and *G*_*T* 3_ for healthy individuals as fixed for AHDS patients. This approach can be interpreted in a physiological sense that the biological maximal activity of D1 (corresponding to *G*_*D*1_ in the model) is the same for healthy individuals and AHDS patients. It is difficult to evaluate whether this approach is reasonable, since most studies only evaluate the *activity* of D1 and not the *maximal activity*. Additionally, the total D1 activity is an extensive parameter, i.e., it depends on the number of expressing cells, approximately on body mass. Therefore, results from cell culture experiments and expression data from biopsies cannot be readily translated to the organismal level.

For example, the activity of D1 inside the thyroid does not change for Mct8 KO mice compared to wild-type littermates (Trajkovic-Arsic et al., 2010a). In turn, no results regarding the maximal activity exist. These observations motivated us to assume a constant maximal activity of D1 and a constant value of *G*_*T* 3_ in our parameter identification, although there are no studies in the literature available examining this fact.

Furthermore, the main challenge of the identification is the observation that the parameters cannot be identified uniquely for healthy individuals. The use of dynamic measurements of the hormone concentrations represents one promising option to tackle this problem. In this case, one could minimize the error of the measured dynamic course of the hormone concentrations and the simulated course of the hormone concentrations. This procedure would be more powerful compared to the current one, in which we minimize the difference between the measured steady-state hormone concentrations and the steady-state hormone concentrations computed by the system, because the informational content is higher in a dynamic course of hormone concentration than in a steady-state value of a hormone concentration.

Furthermore, dynamic hormone measurements would also allow to individualize the mathematical model so that individual AHDS patients can be analyzed. This issue is subject to future research.

## Conclusion

In this paper, we included membrane transporters in the mathematical model of the pituitary-thyroid feedback loop, originally developed by Dietrich (2001); Dietrich et al. (2004); Berberich et al. (2018). The extended model fully replicates the unusual hormone concentrations of AHDS patients and suggests that the retention of *T*_4_ in thyroid cells could fully explain the unusual hormone concentrations of AHDS patients. Currently, the system model represents a generic healthy individual and a generic AHDS patient. One step forward would represent the individualization of the model to account for the different phenotypes of AHDS patients.

## Declaration of Interest

JWD received funding and personal fees by Sanofi-Henning, Hexal AG, Bristol-Myers Squibb, and Pfizer, and is co-owner of the intellectual property rights for the patent “System and Method for Deriving Parameters for Homeostatic Feedback Control of an Individual” (Singapore Institute for Clinical Sciences, Biomedical Sciences Institutes, Application Number 201208940-5, WIPO number WO/2014/088516).

## Funding

This project has received funding from the European Research Council (ERC) under the European Union’s Horizon 2020 research and innovation programme (grant agreement No 948679).

## Contributions

TW drafted the manuscript and performed the calculations/simulations with Matlab/Simulink, under the supervision of MM and with input from JWD. CV derived the different modeling approaches of the membrane transporters and worked on the incorporation of the membrane transporters into the mathematical model, also under the supervision of MM and with input of JWD. Figure 2 was designed by JWD and Julian Berberich (Berberich et al., 2018) and extended by TW. All authors read and approved the manuscript.

## Supplementary Material

The Supplementary Material can be found online.

## Matlab/Simulink Files

The Matlab/Simulink files that were used to perform the simulations and the stability analysis are available online https://doi.org/10.25835/0049176.

## Footnotes

1 In Berberich et al. (2018), these two terms were only present in the differential equation of *T*_3*p*_, because *T*_4,*th*_ was not considered as additional state.

2 The maximal activity regarding the direct *T*_3_ synthesis path is denoted by *G*_*T* 3_, compare Figure 2.

3 We exploit two different studies to parameterize our model, since Groeneweg et al. (2020) only show the *T*_3_ concentrations, but not the *FT*_3_ concentrations.

## Notes

### Competing Interest Statement

Johannes W. Dietrich received funding and personal fees by Sanofi-Henning, Hexal AG, Bristol-Myers Squibb, and Pfizer, and is co-owner of the intellectual property rights for the patent “System and Method for Deriving Parameters for Homeostatic Feedback Control of an Individual” (Singapore Institute for Clinical Sciences, Biomedical Sciences Institutes, Application Number 201208940-5, WIPO number WO/2014/088516).

