## Supplementary Material for "Mathematical Modeling of Thyroid Homeostasis: Implications for the Allan-Herndon-Dudley Syndrome"

#### S1 State of the Art: Mathematical Model of the Pituitary-Thyroid Feedback Loop

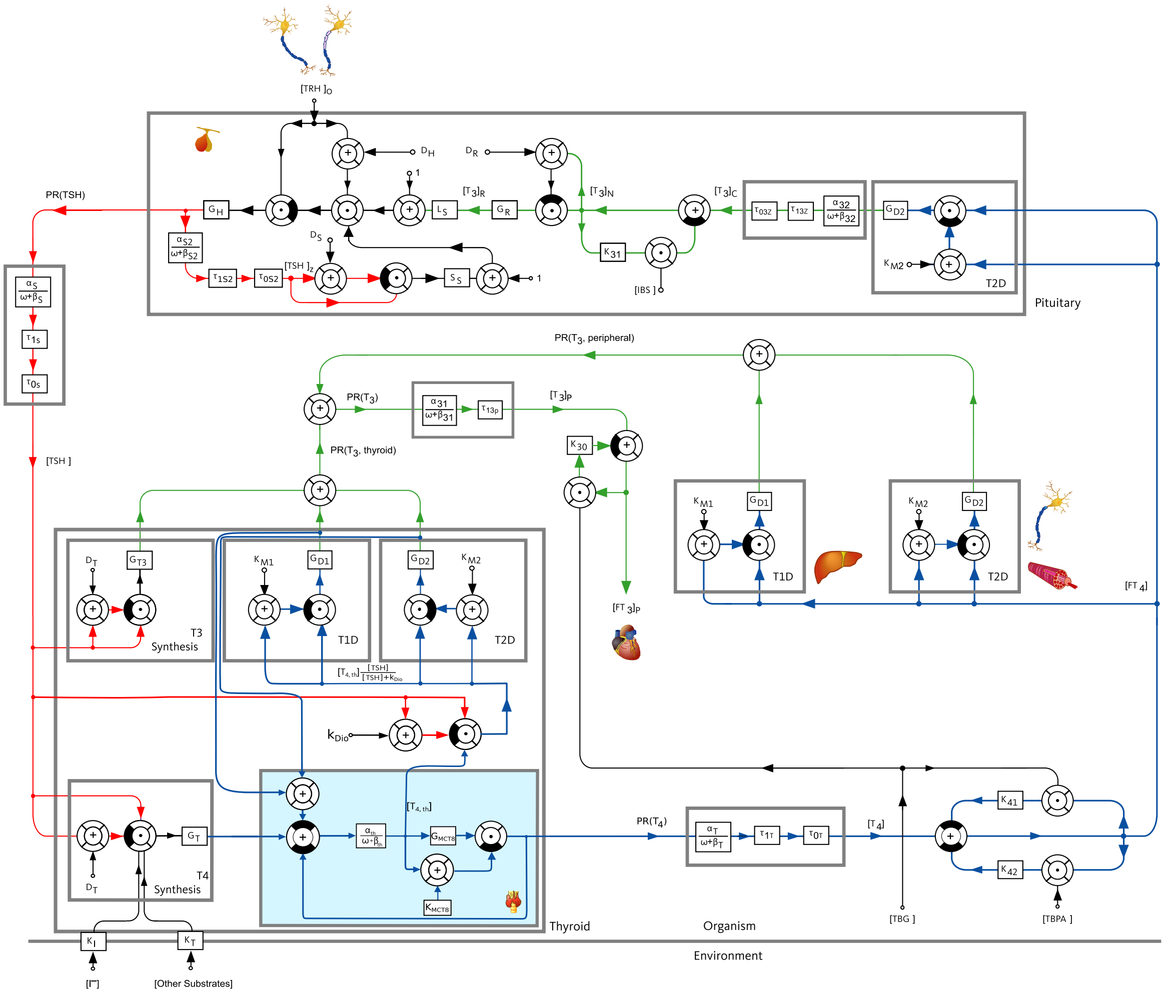

Figure 1: Block diagram of the pituitary-thyroid feedback loop including membrane transporters, extended from Dietrich (2001); Dietrich et al. (2004); Berberich et al. (2018).

As mentioned in the main part of the paper, we want to present in detail the state of the art of the applied mathematical model of the pituitary-thyroid feedback loop. The block diagram of the mathematical model is illustrated in Figure 1.

First, a closer look is taken on the thyroid, which is represented by the grey block in the bottom left corner. In this block, one sees that the $T_4$ synthesis (bottom left corner of the "Thyroid" block) depends on $TSH$, the damping constant of $TSH$ at the thyroid gland ($D_T$), the maximal secretory capacity of the thyroid gland ($G_T$) and the substrate concentrations of thyroglobulin ($K_T$) and iodide ($K_I$). The $T_4$ synthesis is modeled by means of the well known Michaelis-Menten kinetics. The underlying physiological process of the $T_4$ synthesis is of course much more complicated, since it additionally depends on, e.g., the activity of the thyroid peroxidase and sodium/iodide symporter. Nevertheless, a simplification of the underlying physiologic process is necessary in order to keep the model compact.

In the right-hand side of the $T_4$ synthesis block, dilution (denoted by $\alpha_{th}$) and clearance (denoted by $\beta_{th}$) of $T_4$ in thyroid cells take place. Next, the concentration of $T_{4,th}$ is depicted, denoting the $T_4$ concentration in thyroid cells. One part of $T_{4,th}$ is directly transported out of thyroid cells. Here, we denote the maximal activity and the Michaelis-Menten constant of this transport process by $G_{MCT8}$ and $K_{MCT8}$, respectively. The production of $T_3$ in thyroid cells functions through two pathways Berberich et al. (2018). The first pathway is a conversion of $T_{4,th}$ into $T_3$ by means of 5'-deiodinase type I (D1) and 5'-deiodinase type II (D2). This process is modeled via the maximal activities of D1 ($G_{D1}$), of D2 ($G_{D2}$) and the dissociation constants $K_{M1}$, $K_{M2}$, respectively. The second path is a direct synthesis of $T_3$ in thyroid cells, a process, again depending on the concentration of $TSH$, $D_T$ and, additionally, the maximal activity of the $T_3$ synthesis path ($G_{T3}$).

The entrance of $T_4$ in the blood stream is modeled as first order lag element, taking into account dilution ($\alpha_T$) and clearance ($\beta_T$) of $T_4$. Furthermore, a dead time ($\tau_{0T}$) is consider to account for diffusion processes. Additionally, the time constant of the first order lag element ($\tau_{1T}$) is explicitly shown.

Only a small fraction of $T_4$ is available as $FT_4$, since most of it is bound to plasma binding proteins. This phenomenon is taken into account by considering the concentrations of thyroxine-binding globulin ($TBG$), transthyretin ($TBPA$), and their dissociation constants ($K_{41}/K_{42}$) as displayed in the bottom right corner.

The peripheral production of $T_3$ can be seen in the middle of Figure 1. The periphery summarizes the net effects of peripheral organs like the liver or the kidney. This means that we do not explicitly consider the liver and the kidney in the model. We rather summarize their net effects in the periphery. An explicit representation would certainly be helpful in several ways, e.g., to investigate whether the low $T_4$ concentrations of AHDS patients can be explained by an accumulation of $T_4$ in the kidney. However, an explicit representation would go along with new parameters for which numerical values are needed and new states that must be considered. This renders the model even more complex and induces more uncertainty, because the parameters would not necessarily be uniquely identifiable.

In these peripheral organs, D1 and D2 convert $FT_4$ into $FT_3$, a process which is again modeled by means of a Michaelis-Menten kinetics with $G_{D1}$, $G_{D2}$, $K_{M1}$ and $K_{M2}$. This peripheral production is finalized by considering a first-order lag element, taking into account dilution ($\alpha_{31}$) and clearance ($\beta_{31}$). The binding of $T_3$ to $TBG$ and the respective dissociation constant ($K_{41}$) are also visible.

The production of $T_3$ in the pituitary is considered explicitly in the top right-hand corner of Figure 1. It depends on the maximal activity of D2 ($G_{D2}$), the respective dissociation constant ($K_{M2}$), the dilution ($\alpha_{32}$), the clearance factor ($\beta_{32}$) and the dead time ($\tau_{3Z}$). The central $T_3$ ($T_{3z}$) mainly serves as a feedback signal to the pituitary.

Inside the pituitary, which is illustrated in the top of the scheme, many different processes take place. One part of the $T_{3z}$ binds to the intracellular-binding-substrate ($IBS$), which does not serve as a feedback signal. The damping constant $D_R$ models the fact that $T_3$ must bind to specific thyroid hormone receptors in order to influence the concentration of $TSH$. The remaining constants ($G_R$ and $L_S$) stand for the maximum gain of the pituitary receptors in relation to thyroid hormones and a breaking constant, respectively.

Furthermore, the influence of $TRH$ is illustrated in the top of Figure 1. The receptors for $TRH$ at the pituitary are taken into account with a damping constant ($D_H$). The ultra-short feedback loop of $TSH$ on its own secretion (compare Dietrich et al. (2004)) is considered with the respective dilution ($\alpha_{S2}$) and clearance ($\beta_{S2}$) factors. The maximum secretory capacity of the pituitary ($G_H$) is shown. The entrance of $TSH$ into the bloodstream is modeled with a first order lag element, with dilution ($\alpha_S$) and clearance ($\beta_S$).

After having derived the relationships between the different hormone concentrations, one must find the numerical values of all introduced parameters. The parameter values used within this work are listed in Section S6 below. Ideally, we would like to use only human parameters. However, this is not possible for all parameters, consider, e.g., the damping constant of $TRH$ at the pituitary $D_H$, which is not measurable in humans. Therefore, we partially exploit experimentally determined murine numerical parameter values. This results in a model which contains human and murine numerical parameter values. Nevertheless, this approach is meaningful for two reasons: first, there will always be a variation in the exact numerical parameters even for humans only. Second, when applying the mathematical model, we pursue the objective to get insight about the mechanisms of the AHDS, which is also possible when using murine and human numerical parameter values jointly. This is the case since (slightly) different parameter values lead to the same qualitative behavior of hormone concentrations in our model (compare also the sensitivity analysis in Berberich et al. (2018)). On the other hand, an interesting issue for future research could be to fit the model to individual human subjects, e.g., by fitting the unknown parameters to measurement data from an individual subject (similar to what is done for the parameters $G_{D1}$, $G_{T3}$, etc. as described in the main text, compare also Section S2). As discussed in the main text, more (dynamic) measurement data from an individual subject would be needed to this end, i.e., dynamic courses of hormone concentrations obtained through regular/frequent sampling.

#### S2 Constrained Parameter Optimization

Before stating the formal problem of the constrained parameter optimization, it is useful to introduce the system's differential equations for the Michaelis-Menten modeling of the membrane transporters

$$\frac{dT_{4,th}}{dt}(t) = \alpha_{th}\Bigg(G_T\frac{TSH(t)}{TSH(t)+D_T} - G_{MCT8}\frac{T_{4,th}(t)}{K_{MCT8}+T_{4,th}(t)} - G_{D1}\frac{T_{4,th}(t)\frac{TSH(t)}{TSH(t)+k_{Dio}}}{T_{4,th(t)}\frac{TSH(t)}{TSH(t)+k_{Dio}}+K_{M1}} - G_{D2}\frac{T_{4,th}(t)\frac{TSH(t)}{TSH(t)+k_{Dio}}}{T_{4,th}(t)\frac{TSH(t)}{TSH(t)+k_{Dio}}+K_{M2}}\Bigg) - \beta_{th}T_{4,th}(t) \tag{S1}$$

$$\frac{dT_4}{dt}(t) = \alpha_T G_{MCT8}\frac{T_{4,th}(t-\tau_{0T})}{K_{MCT8}+T_{4,th}(t-\tau_{0T})} - \beta_T T_4(t) \tag{S2}$$

$$\frac{dT_{3p}}{dt}(t) = \alpha_{31}\Bigg(G_{D1}\frac{FT_4(t)}{FT_4(t)+K_{M1}} + G_{D2}\frac{FT_4(t)}{FT_4(t)+K_{M2}} + G_{D1}\frac{T_{4,th}(t)\frac{TSH(t)}{TSH(t)+k_{Dio}}}{T_{4,th}(t)\frac{TSH(t)}{TSH(t)+k_{Dio}}+K_{M1}} + G_{D2}\frac{T_{4,th}(t)\frac{TSH(t)}{TSH(t)+k_{Dio}}}{T_{4,th}(t)\frac{TSH(t)}{TSH(t)+k_{Dio}}+K_{M2}} + G_{T3}\frac{TSH(t)}{D_T+TSH(t)}\Bigg) - \beta_{31}T_{3p}(t) \tag{S3}$$

$$\frac{dT_{3c}}{dt}(t) = \alpha_{32}G_{D2}\frac{FT_4(t-\tau_{03Z})}{FT_4(t-\tau_{03Z})+K_{M2}} - \beta_{32}T_{3c}(t) \tag{S4}$$

$$\frac{dTSH}{dt}(t) = \frac{\alpha_S G_H TRH(t-\tau_{0S})}{(TRH(t-\tau_{0S})+D_H)(1+S_S\frac{TSH_z(t-\tau_{0S})}{TSH_z(t-\tau_{0S})+D_S})(1+L_S G_R\frac{T_{3N}(t-\tau_{0S})}{T_{3N}(t-\tau_{0S})+D_R})} - \beta_S TSH(t) \tag{S5}$$

$$\frac{dTSH_z}{dt}(t) = \frac{\alpha_{S2} G_H TRH(t-\tau_{0S2})}{(TRH(t-\tau_{0S2})+D_H)(1+S_S\frac{TSH_z(t-\tau_{0S2})}{TSH_z(t-\tau_{0S2})+D_S})(1+L_S G_R\frac{T_{3N}(t-\tau_{0S2})}{T_{3N}(t-\tau_{0S2})+D_R})} - \beta_{S2} TSH_z(t) \tag{S6}$$

with the relationships

$$FT_3 = T_{3p}\frac{1}{1+K_{30}TBG} \tag{S7}$$

$$FT_4 = T_4\frac{1}{1+K_{41}TBG+K_{42}TBPA} \tag{S8}$$

$$T_{3N} = T_{3c}\frac{1}{1+K_{31}IBS}. \tag{S9}$$

For the linear modeling of the membrane transporters, one must replace the term

$$G_{MCT8}\frac{T_{4,th}}{K_{MCT8}+T_{4,th}} \tag{S10}$$

in equations (S1) and (S2) by

$$k_l T_{4,th}. \tag{S11}$$

As mentioned in the main part of this paper, we perform a constrained parameter optimization in order to identify the parameters $G_{D1}$, $G_{T3}$ and $G_{MCT8}$ or $k_l$, respectively. In other words, we are looking for the configuration of parameters which fits best the given real measured hormone data under the condition that the steady state equations are satisfied.

In formal words, the task is to minimize the objective function $J : \mathbb{R}^7 \to \mathbb{R}$, which is defined by

$$\begin{aligned} J(G_{D1}, G_{T3}, G_{MCT8}, FT_3, FT_4, TSH, T_{4,th}) = &\sum_{n=1}^{81}\left(\frac{FT_{3,i}-FT_3}{\overline{FT_3}}\right)^2 + \sum_{n=1}^{81}\left(\frac{TSH_i-TSH}{\overline{TSH}}\right)^2 \\ &+ \sum_{n=1}^{81}\left(\frac{FT_{4,i}-FT_4}{\overline{FT_4}}\right)^2 \end{aligned} \tag{S12}$$

subject to the nonlinear equality constraints

$$\begin{aligned} 0 = \alpha_{th}\Bigg(&G_T\frac{TSH}{TSH+D_T} - G_{MCT8}\frac{T_{4,th}}{K_{MCT8}+T_{4,th}} - G_{D1}\frac{T_{4,th}\frac{TSH}{TSH+k_{Dio}}}{T_{4,th}\frac{TSH}{TSH+k_{Dio}}+K_{M1}} - G_{D2}\frac{T_{4,th}\frac{TSH}{TSH+k_{Dio}}}{T_{4,th}\frac{TSH}{TSH+k_{Dio}}+K_{M2}}\Bigg) \\ &- \beta_{th}T_{4,th} \end{aligned} \tag{S13}$$

$$0 = FT_4 - \frac{1}{1+K_{41}\,TBG+K_{42}\,TBPA}\frac{\alpha_T}{\beta_T}G_{MCT8}\frac{T_{4,th}}{K_{MCT8}+T_{4,th}} \tag{S14}$$

$$\begin{aligned} 0 = FT_3 - \frac{1}{1+K_{30}TBG}\frac{\alpha_{31}}{\beta_{31}}\Bigg(&G_{D1}\frac{FT_4}{FT_4+K_{M1}} + G_{D2}\frac{FT_4}{FT_4+K_{M2}} + G_{D1}\frac{T_{4,th}\frac{TSH}{TSH+k_{Dio}}}{T_{4,th}\frac{TSH}{TSH+k_{Dio}}+K_{M1}} \\ &+ G_{D2}\frac{T_{4,th}\frac{TSH}{TSH+k_{Dio}}}{T_{4,th}\frac{TSH}{TSH+k_{Dio}}+K_{M2}} + G_{T3}\frac{TSH}{D_T+TSH}\Bigg) \end{aligned} \tag{S15}$$

$$\begin{aligned} 0 = TSH^2\left(G_{MCT8}\frac{T_{4,th}}{K_{MCT8}+T_{4,th}}q_1+q_2\right) + TSH\left(G_{MCT8}\frac{T_{4,th}}{K_{MCT8}+T_{4,th}}q_3+q_4-p_9\right) \\ - G_{MCT8}\frac{T_{4,th}}{K_{MCT8}+T_{4,th}}p_{10} - p_{11} \end{aligned} \tag{S16}$$

and the following bounds

$$G_{D1} \geq 0 \tag{S17}$$

$$G_{T3} \geq 0 \tag{S18}$$

$$G_{MCT8} \geq 0 \tag{S19}$$

$$FT_3 \geq 0 \tag{S20}$$

$$FT_4 \geq 0 \tag{S21}$$

$$TSH \geq 0 \tag{S22}$$

$$T_{4,th} \geq 0. \tag{S23}$$

In (S12), the values $FT_{3,i}$, $TSH_i$ and $FT_{4,i}$ for $i = 1, \ldots, 81$ denote the real hormone measurement data of 81 untreated healthy individuals from Dietrich (2001). The bounds are meaningful given that the parameters $G_{D1}$, $G_{T3}$, $G_{MCT8}$ as well as the hormone concentrations physiologically only make sense when they are positive. We only deal with steady state expressions, therefore, the time dependence of the variables is neglected. Note that we plugged in the steady state expressions of the differential equations of $T_{3c}$ and of $TSH_z$ into the steady state expression of $TSH$. This leads to condition (S16). The detailed steps how to reach (S16), which are cumbersome but straightforward, are shown in Section S5 for completeness.

One challenge of this approach is that the order of magnitude of the constraints is very different. The value of $FT_3$ is of order $10^{-12}$, whereas the order of $TSH$ is 1. In order to obtain a numerically well conditioned problem, we hence perform the following state transformation

$$\begin{pmatrix} \widetilde{T_{4,th}} \\ \widetilde{T_4} \\ \widetilde{T_{3p}} \\ \widetilde{T_{3c}} \\ \widetilde{TSH} \\ \widetilde{TSH_z} \end{pmatrix} = \mathbf{z} = \mathbf{T}\mathbf{x} = \mathbf{T} \begin{pmatrix} T_{4,th} \\ T_4 \\ T_{3p} \\ T_{3c} \\ TSH \\ TSH_z \end{pmatrix}, \tag{S24}$$

with

$$\mathbf{T} = \begin{pmatrix} 10^{12} & 0 & 0 & 0 & 0 & 0 \\ 0 & 10^{11} & 0 & 0 & 0 & 0 \\ 0 & 0 & 10^{12} & 0 & 0 & 0 \\ 0 & 0 & 0 & 10^{8} & 0 & 0 \\ 0 & 0 & 0 & 0 & 1 & 0 \\ 0 & 0 & 0 & 0 & 0 & 1 \end{pmatrix}. \tag{S25}$$

This transformation matrix leads to (transformed) hormone concentrations in the identification ($FT_3$, $FT_4$, $T_{4,th}$ and $TSH$) that are in the order of magnitude of $10^0$. Additionally, we set $\widetilde{G_{D1}} = 10^8 G_{D1}$, $\widetilde{G_{T3}} = 10^{14} G_{T3}$ and $\widetilde{G_{MCT8}} = 10^6 G_{MCT8}$ such that all variables in the optimization problem (S12)-(S23) are in the same order of magnitude.

The dynamics of the transformed system are

$$\dot{\mathbf{z}} = \mathbf{T}\dot{\mathbf{x}} = \mathbf{T}\mathbf{f}(\mathbf{x}) = \mathbf{T}\mathbf{f}(\mathbf{T}^{-1}\mathbf{z}), \tag{S26}$$

where $\mathbf{f}(\mathbf{x})$ denotes the right hand side of the differential equations (S1)-(S6) and $x$, $z$ as defined in (S24). When the mentioned values are plugged in, the expression

$$\dot{\mathbf{z}} = \begin{pmatrix} 10^{12} & 0 & 0 & 0 & 0 & 0 \\ 0 & 10^{11} & 0 & 0 & 0 & 0 \\ 0 & 0 & 10^{12} & 0 & 0 & 0 \\ 0 & 0 & 0 & 10^8 & 0 & 0 \\ 0 & 0 & 0 & 0 & 1 & 0 \\ 0 & 0 & 0 & 0 & 0 & 1 \end{pmatrix} \mathbf{f} \begin{pmatrix} 10^{-12}\widetilde{T_{4,th}} \\ 10^{-11}\widetilde{T_4} \\ 10^{-12}\widetilde{T_{3p}} \\ 10^{-8}\widetilde{T_{3c}} \\ \widetilde{TSH} \\ \widetilde{TSH_z} \end{pmatrix} \tag{S27}$$

is obtained.

The transformed differential equations are

$$\frac{d\widetilde{T_{4,th}}}{dt} = 10^{12}\left(\alpha_{th}\left(G_T\frac{\widetilde{TSH}}{\widetilde{TSH}+D_T} - 10^{-6}\widetilde{G_{MCT8}}\frac{10^{-12}\widetilde{T_{4,th}}}{K_{MCT8}+10^{-12}\widetilde{T_{4,th}}} - 10^{-8}\widetilde{G_{D1}}\frac{10^{-12}\widetilde{T_{4,th}}\frac{\widetilde{TSH}}{\widetilde{TSH}+k_{Dio}}}{10^{-12}\widetilde{T_{4,th}}\frac{\widetilde{TSH}}{\widetilde{TSH}+k_{Dio}}+K_{M1}}\right.\right.$$
$$\left.\left. - G_{D2}\frac{10^{-12}\widetilde{T_{4,th}}\frac{\widetilde{TSH}}{\widetilde{TSH}+k_{Dio}}}{10^{-12}\widetilde{T_{4,th}}\frac{\widetilde{TSH}}{\widetilde{TSH}+k_{Dio}}+K_{M2}}\right) - \beta_{th}T_{4,th}\right) \tag{S28}$$

$$\frac{d\widetilde{T_4}}{dt} = 10^{11}\left(\alpha_T\,10^{-6}\widetilde{G_{MCT8}}\frac{10^{-12}\widetilde{T_{4,th}}}{K_{MCT8}+10^{-12}\widetilde{T_{4,th}}} - \beta_T 10^{-11}\widetilde{T_4}\right) \tag{S29}$$

$$\frac{d\widetilde{T_{3p}}}{dt} = 10^{12}\left(\alpha_{31}\left(10^{-8}\widetilde{G_{D1}}\frac{10^{-11}\widetilde{FT_4}}{10^{-11}\widetilde{FT_4}+K_{M1}} + G_{D2}\frac{10^{-11}\widetilde{FT_4}}{10^{-11}\widetilde{FT_4}+K_{M2}}\right.\right.$$
$$+ 10^{-8}\widetilde{G_{D1}}\frac{10^{-12}\widetilde{T_{4,th}}\frac{\widetilde{TSH}}{\widetilde{TSH}+k_{Dio}}}{10^{-12}\widetilde{T_{4,th}}\frac{\widetilde{TSH}}{\widetilde{TSH}+k_{Dio}}+K_{M1}} + G_{D2}\frac{10^{-12}\widetilde{T_{4,th}}\frac{\widetilde{TSH}}{\widetilde{TSH}+k_{Dio}}}{10^{-12}\widetilde{T_{4,th}}\frac{\widetilde{TSH}}{\widetilde{TSH}+k_{Dio}}+K_{M2}}$$
$$\left.\left. + 10^{-14}\widetilde{G_{T3}}\frac{\widetilde{TSH}}{D_T+\widetilde{TSH}}\right) - \beta_{31}10^{-12}\widetilde{T_{3p}}\right) \tag{S30}$$

$$\frac{d\widetilde{T_{3c}}}{dt} = 10^{8}\left(\alpha_{32}G_{D2}\frac{10^{-11}\widetilde{FT_4}}{10^{-11}\widetilde{FT_4}+K_{M2}} - \beta_{32}10^{-8}\widetilde{T_{3c}}\right) \tag{S31}$$

$$\frac{d\widetilde{TSH}}{dt} = \frac{\alpha_S G_H TRH}{(TRH+D_H)\left(1+S_S\frac{\widetilde{TSH_z}}{\widetilde{TSH_z}+D_S}\right)\left(1+L_S G_R\frac{10^{-8}\widetilde{T_{3N}}}{10^{-8}\widetilde{T_{3N}}+D_R}\right)} - \beta_S\widetilde{TSH} \tag{S32}$$

$$\frac{d\widetilde{TSH_z}}{dt} = \frac{\alpha_{S2} G_H TRH}{(TRH+D_H)\left(1+S_S\frac{\widetilde{TSH_z}}{\widetilde{TSH_z}+D_S}\right)\left(1+L_S G_R\frac{10^{-8}\widetilde{T_{3N}}}{10^{-8}\widetilde{T_{3N}}+D_R}\right)} - \beta_{S2}\widetilde{TSH_z}\,. \tag{S33}$$

Furthermore, the relationships of the hormone concentrations

$$FT_3 = 10^{-12}\widetilde{T_{3p}}\frac{1}{1+K_{30}TBG} = 10^{-12}\widetilde{FT_3} \tag{S34}$$
$$\widetilde{FT_3} = \widetilde{T_{3p}}\frac{1}{1+K_{30}TBG} \tag{S35}$$

$$FT_4 = 10^{-11}\widetilde{T_4}\frac{1}{1+K_{41}\,TBG+K_{42}\,TBPA} = 10^{-11}\widetilde{FT_4} \tag{S36}$$
$$\widetilde{FT_4} = \widetilde{T_4}\frac{1}{1+K_{41}\,TBG+K_{42}\,TBPA} \tag{S37}$$

$$T_{3N} = 10^{-8}\widetilde{T_{3c}}\frac{1}{1+K_{31}IBS} = 10^{-8}\widetilde{T_{3N}} \tag{S38}$$
$$\widetilde{T_{3N}} = \widetilde{T_{3c}}\frac{1}{1+K_{31}IBS} \tag{S39}$$

need to be considered. Using this state transformation, all variables are in the same order of magnitude. We can now formulate the following transformed optimization problem, which is numerically better conditioned. The objective function $J : \mathbb{R}^7 \to \mathbb{R}$ defined by

$$J(\widetilde{G_{D1}}, \widetilde{G_{T3}}, \widetilde{G_{MCT8}}, \widetilde{FT_3}, \widetilde{FT_4}, \widetilde{TSH}, \widetilde{T_{4,th}}) = \sum_{n=1}^{81} \left( \frac{FT_{3,i} - 10^{-12}\widetilde{FT_3}}{\overline{FT_3}} \right)^2 + \sum_{n=1}^{81} \left( \frac{TSH_i - \widetilde{TSH}}{\overline{TSH}} \right)^2 + \sum_{n=1}^{81} \left( \frac{FT_{4,i} - 10^{-11}\widetilde{FT_4}}{\overline{FT_4}} \right)^2 \tag{S40}$$

must be minimized. The problem is subject to the nonlinear constraints

$$0 = 10^{12}\Bigg(\alpha_{th}\Bigg(G_T \frac{\widetilde{TSH}}{\widetilde{TSH} + D_T} - 10^{-6}\widetilde{G_{MCT8}} \frac{10^{-12}\widetilde{T_{4,th}}}{K_{MCT8} + 10^{-12}\widetilde{T_{4,th}}} - 10^{-8}\widetilde{G_{D1}} \frac{10^{-12}\widetilde{T_{4,th}}\frac{\widetilde{TSH}}{\widetilde{TSH}+k_{Dio}}}{10^{-12}\widetilde{T_{4,th}}\frac{\widetilde{TSH}}{\widetilde{TSH}+k_{Dio}} + K_{M1}} - G_{D2} \frac{10^{-12}\widetilde{T_{4,th}}\frac{\widetilde{TSH}}{\widetilde{TSH}+k_{Dio}}}{10^{-12}\widetilde{T_{4,th}}\frac{\widetilde{TSH}}{\widetilde{TSH}+k_{Dio}} + K_{M2}}\Bigg) - \beta_{th} T_{4,th}\Bigg) \tag{S41}$$

$$0 = \widetilde{FT_4} - \frac{1}{1 + K_{41}\,TBG + K_{42}\,TBPA} \frac{\alpha_T}{\beta_T} \frac{1}{10^{-11}} 10^{-6}\widetilde{G_{MCT8}} \frac{10^{-12}\widetilde{T_{4,th}}}{K_{MCT8} + 10^{-12}\widetilde{T_{4,th}}} \tag{S42}$$

$$0 = \widetilde{FT_3} - \frac{1}{1 + K_{30}TBG} \frac{\alpha_{31}}{\beta_{31}} \frac{1}{10^{-12}} \Bigg(10^{-8}\widetilde{G_{D1}} \frac{10^{-11}\widetilde{FT_4}}{10^{-11}\widetilde{FT_4} + K_{M1}} + G_{D2} \frac{10^{-11}\widetilde{FT_4}}{10^{-11}\widetilde{FT_4} + K_{M2}} + 10^{-8}\widetilde{G_{D1}} \frac{10^{-12}\widetilde{T_{4,th}}\frac{\widetilde{TSH}}{\widetilde{TSH}+k_{Dio}}}{10^{-12}\widetilde{T_{4,th}}\frac{\widetilde{TSH}}{\widetilde{TSH}+k_{Dio}} + K_{M1}} + G_{D2} \frac{10^{-12}\widetilde{T_{4,th}}\frac{\widetilde{TSH}}{\widetilde{TSH}+k_{Dio}}}{10^{-12}\widetilde{T_{4,th}}\frac{\widetilde{TSH}}{\widetilde{TSH}+k_{Dio}} + K_{M2}} + 10^{-14}\widetilde{G_{T3}} \frac{\widetilde{TSH}}{D_T + \widetilde{TSH}}\Bigg) \tag{S43}$$

$$0 = \widetilde{TSH}^2 (10^{-6}\widetilde{G_{MCT8}} \frac{10^{-12}\widetilde{T_{4,th}}}{K_{MCT8} + 10^{-12}\widetilde{T_{4,th}}} q_1 + q_2) + \widetilde{TSH}(10^{-6}\widetilde{G_{MCT8}} \frac{10^{-12}\widetilde{T_{4,th}}}{K_{MCT8} + 10^{-12}\widetilde{T_{4,th}}} q_3 + q_4 - p_9) - 10^{-6}\widetilde{G_{MCT8}} \frac{10^{-12}\widetilde{T_{4,th}}}{K_{MCT8} + 10^{-12}\widetilde{T_{4,th}}} p_{10} - p_{11} \tag{S44}$$

and the following bounds

$$\widetilde{G_{D1}} \geq 0 \tag{S45}$$

$$\widetilde{G_{T3}} \geq 0 \tag{S46}$$

$$\widetilde{G_{MCT8}} \geq 0 \tag{S47}$$

$$\widetilde{FT_3} \geq 0 \tag{S48}$$

$$\widetilde{FT_4} \geq 0 \tag{S49}$$

$$\widetilde{TSH} \geq 0 \tag{S50}$$

$$\widetilde{T_{4,th}} \geq 0. \tag{S51}$$

Analogous to (S16), constraint (S44) follows from plugging in the steady-state equations of $\widetilde{T_{3c}}$ (S31) and $\widetilde{TSH_z}$ (S33) into the steady-state equation of $\widetilde{TSH}$ (S32). As all variables are in the same order of magnitude, the problem is now much better conditioned.

The optimization problem (S40)-(S51) can be used for the parameter identification of healthy individuals. The parameter identification of AHDS patients can be set up in a similar fashion. For AHDS patients, we fit the unknown parameters to the mean values of the measured hormone concentrations from Groeneweg et al. (2020) and Schwartz et al. (2005), compare Section Methods of the main paper. Therefore, the sums in the cost function only contain one element, i.e., the mean value of the corresponding hormone concentration. Furthermore, as discussed in Section Methods of the main paper, for AHDS patients we keep the values of $G_{D1}$ and $G_{T3}$ fixed (which means that they are not treated as optimization variables) and only identify $G_{MCT8}$.

Regarding the parameter identification of the linear modeling of the membrane transporters, the change of the system's dynamics described by equations (S10) and (S11) must be performed. Then, the parameter identification works analogously.

### S3 Results Constrained Parameter Optimization

This section is dedicated to the detailed results of the constrained parameter optimization. These results were obtained using an Intel(R) Core(TM) i7-10875H CPU @ 2.30GHz (16 CPUs) processor with 16 GB RAM and MATLAB®, 9.10.0.1684407 (R2021a). We used the built-in MATLAB solver *fmincon*.

Especially, we want to analyze the observation that the parameters are not uniquely identifiable for healthy individuals in more detail. First of all, one can take a closer look on Table S1. In this Table, different initial guesses and the corresponding final values of the optimization (corresponding to local minima of the optimization problem (S40)-(S51)) are illustrated. Compared to case "Healthy 1", the changed initial guesses of the constrained parameter optimization are always shown in italics. When comparing for example case "Healthy

Table S1: Results of the parameter identification for the Michaelis-Menten modeling of the membrane transporters with the constrained parameter optimization approach

| Case | | Parameters to be Identified | | | Corresponding Hormone Concentrations | | | | Further Characteristics | | |
|---|---|---|---|---|---|---|---|---|---|---|---|
| | | $G_{D1}$ in $10^{-8}\frac{\text{mol}}{\text{s}}$ | $G_{T3}$ in $10^{-14}\frac{\text{mol}}{\text{s}}$ | $G_{MCT8}$ in $10^{-6}\frac{\text{mol}}{\text{s}}$ | $FT_3$ in $10^{-12}\frac{\text{mol}}{\text{l}}$ | $FT_4$ in $10^{-11}\frac{\text{mol}}{\text{l}}$ | $TSH$ in $\frac{\text{mIU}}{\text{l}}$ | $T_{4,th}$ in $10^{-12}\frac{\text{mol}}{\text{l}}$ | Periphery % | Thyroid % | Costs |
| Healthy 1 | Initial | 2.2 | 21 | 4.7 | 4.50 | 1.41 | 1.40 | 1.14 | | | |
| | Final | 1.98 | 20.90 | 1.94 | 4.50 | 1.69 | 1.86 | 3.10 | ≈ 80 | ≈ 20 | 34.50 |
| Healthy 2 | Initial | 2.2 | *10* | 4.7 | 4.50 | 1.41 | 1.40 | 1.14 | | | |
| | Final | 2.11 | 9.87 | 2.07 | 4.50 | 1.69 | 1.86 | 2.91 | ≈ 86 | ≈ 14 | 34.50 |
| Healthy 3 | Initial | *4* | 21 | 4.7 | 4.50 | 1.41 | 1.40 | 1.14 | | | |
| | Final | 1.98 | 20.89 | 1.94 | 4.50 | 1.69 | 1.86 | 3.10 | ≈ 80 | ≈ 20 | 34.50 |
| Healthy 4 | Initial | 2.2 | 21 | *10* | 4.50 | 1.41 | 1.4 | 1.14 | | | |
| | Final | 1.98 | 20.43 | 1.95 | 4.50 | 1.69 | 1.86 | 3.10 | ≈ 80 | ≈ 20 | 34.50 |
| Healthy 5 | Initial | 2.2 | *30* | 4.7 | 4.50 | 1.41 | 1.40 | 1.14 | | | |
| | Final | 1.87 | 29.91 | 1.84 | 4.50 | 1.69 | 1.86 | 3.28 | ≈ 76 | ≈ 24 | 34.50 |
| AHDS | Initial | — | — | 1.94 | 8.14 | 0.95 | 2.97 | 3.10 | | | |
| | Final | — | — | 0.09 | 7.83 | 0.88 | 2.67 | 34.68 | ≈ 24 | ≈ 76 | 0.0170 |

Annotations: the further characteristics evaluate the percentage of $T_3$, which is produced in the periphery and in the thyroid.

1" and case "Healthy 2", it becomes apparent that different final values of $G_{D1}$, $G_{T3}$ and $G_{MCT8}$ lead to the same values of $FT_3$, $FT_4$ and $TSH$. Note that the costs of these two cases are then identical as well.

In the following, we discuss how this observation can be explained mathematically. We focus on the formulation of constrained parameter optimization without the mentioned transformation, since it might be easier to follow for the reader. In the cost function of the constrained parameter optimization (S12), there are three different terms, one related to $FT_3$, one related to $FT_4$ and one related to $TSH$. The constraints of $FT_4$ (S14) and $TSH$ (S16) only depend on $G_{MCT8}$ and $T_{4,th}$. One can manipulate the respective constraints and obtain the expressions

$$FT_4(G_{MCT8}, T_{4,th}) = \frac{1}{1 + K_{41}\,TBG + K_{42}\,TBPA}\frac{\alpha_T}{\beta_T}G_{MCT8}\frac{T_{4,th}}{K_{MCT8} + T_{4,th}} \tag{S52}$$

$$TSH(G_{MCT8}, T_{4,th}) = -\frac{G_{MCT8}\frac{T_{4,th}}{K_{MCT8}+T_{4,th}}q_3 + q_4 + p_9}{2(G_{MCT8}\frac{T_{4,th}}{K_{MCT8}+T_{4,th}}q_1 + q_2)} + \sqrt{\left(\frac{G_{MCT8}\frac{T_{4,th}}{K_{MCT8}+T_{4,th}}q_3 + q_4 + p_9}{2(G_{MCT8}\frac{T_{4,th}}{K_{MCT8}+T_{4,th}}q_1 + q_2)}\right)^2 + \frac{G_{MCT8}\frac{T_{4,th}}{K_{MCT8}+T_{4,th}}p_{10} + p_{11}}{G_{MCT8}\frac{T_{4,th}}{K_{MCT8}+T_{4,th}}q_1 + q_2}}, \tag{S53}$$

Table S2: Results of the parameter identification for the linear modeling of the membrane transporters with the constrained parameter optimization approach

| Case | | Parameters to be Identified | | | Corresponding Hormone Concentrations | | | | Further Characteristics | | |
|---|---|---|---|---|---|---|---|---|---|---|---|
| | | $G_{D1}$ in $10^{-8}\frac{\text{mol}}{\text{s}}$ | $G_{T3}$ in $10^{-14}\frac{\text{mol}}{\text{s}}$ | $k_l$ in $\frac{1}{\text{s}}$ | $FT_3$ in $10^{-12}\frac{\text{mol}}{\text{l}}$ | $FT_4$ in $10^{-11}\frac{\text{mol}}{\text{l}}$ | $TSH$ in $\frac{\text{mIU}}{\text{l}}$ | $T_{4,th}$ in $10^{-12}\frac{\text{mol}}{\text{l}}$ | Periphery % | Thyroid % | Costs |
| Healthy 1 | Initial | 2.2 | 21 | 1 | 4.50 | 1.41 | 1.40 | 1.14 | | | |
| | Final | 1.98 | 20.97 | 0.4130 | 4.50 | 1.69 | 1.86 | 3.11 | ≈ 80 | ≈ 20 | 34.50 |
| Healthy 2 | Initial | 2.2 | *10* | 1 | 4.50 | 1.41 | 1.40 | 1.14 | | | |
| | Final | 2.11 | 9.97 | 0.4404 | 4.50 | 1.69 | 1.86 | 2.91 | ≈ 86 | ≈ 14 | 34.50 |
| Healthy 3 | Initial | *4* | 21 | 1 | 4.50 | 1.41 | 1.40 | 1.14 | | | |
| | Final | 1.98 | 20.96 | 0.4130 | 4.50 | 1.69 | 1.86 | 3.11 | ≈ 80 | ≈ 20 | 34.50 |
| Healthy 4 | Initial | 2.2 | 21 | *2* | 4.50 | 1.41 | 1.40 | 1.14 | | | |
| | Final | 1.98 | 20.95 | 0.4130 | 4.50 | 1.69 | 1.86 | 3.11 | ≈ 80 | ≈ 20 | 34.50 |
| Healthy 5 | Initial | 2.2 | *30* | 1 | 4.50 | 1.41 | 1.40 | 1.14 | | | |
| | Final | 1.87 | 29.95 | 0.3906 | 4.50 | 1.69 | 1.86 | 3.28 | ≈ 75 | ≈ 24 | 34.50 |
| AHDS | Initial | — | — | 0.4130 | 8.14 | 0.95 | 2.97 | 3.11 | | | |
| | Final | — | — | 0.0192 | 7.83 | 0.88 | 2.67 | 34.69 | ≈ 24 | ≈ 76 | 0.0170 |

Annotations: the further characteristics evaluate the percentage of $T_3$, which is produced in the periphery and in the thyroid.

where only one solution of the quadratic equation of $TSH$ is positive for a meaningful range of possible $T_{4,th}$ and $G_{MCT8}$ values[1]. Since $K_{MCT8} >> T_{4,th}$, one can approximate the nonlinear expression

$$G_{MCT8}\frac{T_{4,th}}{K_{MCT8}+T_{4,th}} \tag{S54}$$

by

$$\frac{G_{MCT8}}{K_{MCT8}}T_{4,th}. \tag{S55}$$

Because the value of $K_{MCT8}$ is fixed, it follows that the optimal values of $FT_4$ and $TSH$ (approximately) only depend on the product of $G_{MCT8}$ and $T_{4,th}$. The isolated values of $FT_4$ and $TSH$ are not uniquely determined by the remaining constraints of the optimization problem (S13) and (S15), because (S13) depends linearly on $G_{D1}$ and (S15) depends linearly on $G_{T3}$. For any positive values of $T_{4,th}$ and $G_{MCT8}$, one can adapt the value of $G_{D1}$ in order to fulfill (S13) and $G_{T3}$ in order to fulfill (S15). This means that an infinite number of combinations of $G_{MCT8}$ and $T_{4,th}$ exist which result in the same product and hence (approximately) the same $FT_4$ and $TSH$ values. Note that in all documented cases in Table S1, the product of $G_{MCT8}$ and $T_{4,th}$ indeed remains approximately the same, namely $\approx 5.9\cdot 10^{-18}\frac{\text{mol}^2}{\text{sl}}$, but that their individual values vary.

These different values impact the result of the other parameters, namely $G_{D1}$ and $G_{T3}$. For given $T_{4,th}$,

[1] The meaning of all variables is explained in Sections S5 and S6

$TSH$ and $G_{MCT8}$, it follows that $G_{D1}$ must be such that constraint (S13) is satisfied, resulting in different $G_{D1}$ values for varying $T_{4,th}$ and $G_{MCT8}$ values. By means of the last condition (S15), one can explain the last varying parameter, namely $G_{T3}$. In this last equality constraint, the optimal value of $FT_3$ is determined by the respective term in the cost function, but as mentioned above, the values of $T_{4,th}$ and $G_{D1}$ can vary. This leads to a variation of the last unknown parameter, namely $G_{T3}$, which must be such that (S15) is satisfied.

Interestingly, the optimal value of $FT_3$ is even equal to the mean measured hormone concentration. This is due to the observation that the constraint related to $FT_3$ can always be fulfilled by the adaptation of $G_{T3}$ and thus the optimal value of $FT_3$ just corresponds to the mean measured hormone concentration. This is not the case for the optimal values of $FT_4$ and $TSH$, which are different from the mean measured hormone concentrations.

Regarding the variations of the linear modeling of the membrane transporters, the explanation is completely analogous. The detailed results are illustrated in Table S2.

### S4 Stability Analysis

As mentioned in the main part of this paper, we performed a local stability analysis. To this end, we introduce Lyapunov's indirect method. Then, we document and discuss the results from a physiological point of view.

#### Lyapunov's Indirect Method

Lyapunov's indirect method can be formulated as follows (Khalil, 2002): let $x = 0$ be an equilibrium point/hormone concentration for the nonlinear system

$$\dot{\mathbf{x}} = \mathbf{f}(\mathbf{x}) \tag{S56}$$

where $\mathbf{f} : D \mapsto \mathbb{R}^n$ is continuously differentiable and D is a neighborhood of the origin. Let

$$\mathbf{A} = \left.\frac{\partial \mathbf{f}(\mathbf{x})}{\partial \mathbf{x}}\right|_{\mathbf{x}=0}, \tag{S57}$$

then

1. the origin is asymptotically stable if Re $\lambda_i < 0$ for all eigenvalues of $\mathbf{A}$.
2. The origin is unstable if Re $\lambda_i > 0$ for one or more of the eigenvalues of $\mathbf{A}$.

As Lyapunov's indirect method is only a criterion for local asymptotic stability, no statement is possible regarding the region of attraction of the equilibrium point. The region of attraction of an equilibrium point is the set of all initial states (here: hormone concentrations), for which the solution of the nonlinear system (S57) asymptotically converges to the equilibrium point.

Table S3: Results of the application of Lyapunov's indirect method.

| | M.-M. Modeling of Membrane Transporters | | Linear Modeling of Membrane Transporters | |
|---|---|---|---|---|
| Eigenvalue | Healthy Individuals | AHDS Patients | Healthy Individuals | AHDS Patients |
| $\lambda_1$ | $-8.00 \cdot 10^{-6}$ | $-8.00 \cdot 10^{-6}$ | $-8.00 \cdot 10^{-6}$ | $-8.00 \cdot 10^{-6}$ |
| $\lambda_2$ | $-2.47 \cdot 10^{2}$ | $-2.52 \cdot 10^{2}$ | $-2.47 \cdot 10^{2}$ | $-2.52 \cdot 10^{2}$ |
| $\lambda_3$ | $-8.30 \cdot 10^{-4}$ | $-8.30 \cdot 10^{-4}$ | $-8.30 \cdot 10^{-4}$ | $-8.30 \cdot 10^{-4}$ |
| $\lambda_4$ | $-1.45 \cdot 10^{-6}$ | $-1.31 \cdot 10^{-6}$ | $-1.45 \cdot 10^{-6}$ | $-1.31 \cdot 10^{-6}$ |
| $\lambda_5$ | $-2.30 \cdot 10^{-4}$ | $-2.30 \cdot 10^{-4}$ | $-2.30 \cdot 10^{-4}$ | $-2.30 \cdot 10^{-4}$ |
| $\lambda_6$ | $-109.72$ | $-12.00$ | $-109.67$ | $-11.99$ |

Annotations: the abbreviation M.M. stands for Michaelis-Menten.

#### Results Lyapunov's Indirect Method

The stability analysis concerns the equilibrium hormone concentrations documented in Tables 1 and 2 of the main part, which were obtained using a constant concentration of $TRH$. The application of Lyapunov's indirect method leads to the results illustrated in Table S3. One can see that the eigenvalues are negative for all cases. Consequently, the computed equilibrium hormone concentrations are all locally asymptotically stable.

#### Discussion Stability Analysis

After having documented these (rather technical) results, we can focus on their medical interpretation.

Local asymptotic stability means that if the hormone concentrations have values different from their equilibrium hormone concentrations, which are still in the region of attraction, the hormone concentrations will converge to their equilibrium hormone concentrations. Unfortunately, with the proposed method it is impossible to state how large the region of attraction is. Hence, it is impossible to quantify the maximal deviation of the hormone concentrations for which they converge to their equilibrium hormone concentrations.

This results holds true for healthy individuals and for AHDS patients. This is particularly interesting since AHDS patients have a highly perturbed pituitary-thyroid feedback loop. From a systems theoretic point of view, their equilibrium points are still locally asymptotically stable.

In the future, it would be highly valuable to determine the mentioned region of attraction. A possible method that goes along with a quantification of the region of attraction is Lyapunov's direct method (Khalil, 2002). However, its application is highly challenging if the differential equations are complex as it is the case here.

#### S5 Derivation of the Steady State Equation of TSH

Here the step-by-step solution to derive the steady-state equation of $TSH$ in dependence of $T_{4,th}$ is presented for the Michaelis-Menten modeling of the membrane transporters.

The steady-state equation of $T_4$ is

$$T_4 = \frac{\alpha_T}{\beta_T} G_{MCT8} \frac{T_{4,th}}{K_{MCT8} + T_{4,th}}, \tag{S58}$$

which follows from setting (S2) to zero and solving the equation for $T_4$. This steady-state equation can be plugged into the relationship of $FT_4$ (S8), thus leading to

$$FT_4 = \frac{\alpha_T}{\beta_T} G_{MCT8} \frac{T_{4,th}}{K_{MCT8} + T_{4,th}} \frac{1}{1 + K_{41}TBG + K_{42}TBPA}. \tag{S59}$$

With $b_1 = 1/(1 + K_{41}TBG + K_{42}TBPA)$, equation (S59) can be written as

$$FT_4 = \frac{\alpha_T}{\beta_T} G_{MCT8} \frac{T_{4,th}}{K_{MCT8} + T_{4,th}} b_1. \tag{S60}$$

This can be used for the steady state equation of $T_{3c}$, which can again be computed by setting (S4) to zero and solving for $T_{3c}$. When the relation of $FT_4$ (S60) is plugged into this steady-state equation, one gets

$$T_{3c} = a_3 \frac{\frac{\alpha_T}{\beta_T} G_{MCT8} \frac{T_{4,th}}{K_{MCT8}+T_{4,th}} b_1}{\frac{\alpha_T}{\beta_T} G_{MCT8} \frac{T_{4,th}}{K_{MCT8}+T_{4,th}} b_1 + K_{M2}} \tag{S61}$$

with $a_3 = \alpha_{32} G_{D2}/\beta_{32}$. By defining $p_1 = \alpha_T b_1/\beta_T$, equation (S61) simplifies to

$$T_{3c} = a_3 \frac{p_1 G_{MCT8} \frac{T_{4,th}}{K_{MCT8}+T_{4,th}}}{p_1 G_{MCT8} \frac{T_{4,th}}{K_{MCT8}+T_{4,th}} + K_{M2}}. \tag{S62}$$

Next, the expression of $T_{3N}$ (S9) is used. With $b_3 = 1/(1 + K_{31}IBS)$ and expression (S62), $T_{3N}$ is

$$T_{3N} = b_3 T_{3c} \tag{S63}$$

$$= b_3 a_3 \frac{p_1 G_{MCT8} \frac{T_{4,th}}{K_{MCT8}+T_{4,th}}}{p_1 G_{MCT8} \frac{T_{4,th}}{K_{MCT8}+T_{4,th}} + K_{M2}}. \tag{S64}$$

The term $G_R T_{3N}/(T_{3N} + D_R)$ is needed for the steady-state computation of $TSH$. With the definition of $p_2 = a_3 b_3 p_1$ and $p_3 = p_2 + D_R p_1$, it becomes

$$G_R \frac{T_{3N}}{T_{3N}+D_R} = G_R \frac{b_3 a_3 \frac{p_1 G_{MCT8} \frac{T_{4,th}}{K_{MCT8}+T_{4,th}}}{p_1 G_{MCT8} \frac{T_{4,th}}{K_{MCT8}+T_{4,th}} + K_{M2}}}{b_3 a_3 \frac{p_1 G_{MCT8} \frac{T_{4,th}}{K_{MCT8}+T_{4,th}}}{p_1 G_{MCT8} \frac{T_{4,th}}{K_{MCT8}+T_{4,th}} + K_{M2}} + D_R} \tag{S65}$$

$$= G_R \frac{b_3 a_3 p_1 G_{MCT8} \frac{T_{4,th}}{K_{MCT8}+T_{4,th}}}{b_3 a_3 p_1 G_{MCT8} \frac{T_{4,th}}{K_{MCT8}+T_{4,th}} + D_R (p_1 G_{MCT8} \frac{T_{4,th}}{K_{MCT8}+T_{4,th}} + K_{M2})} \tag{S66}$$

$$= G_R \frac{p_2 G_{MCT8} \frac{T_{4,th}}{K_{MCT8}+T_{4,th}}}{p_3 G_{MCT8} \frac{T_{4,th}}{K_{MCT8}+T_{4,th}} + D_R K_{M2}}. \tag{S67}$$

Now the steady-state value of $TSH$ can be computed, which again follows from setting (S5) to zero and solving for $TSH$. With the following relationships

$$p_4 = G_H \frac{\alpha_S}{\beta_S} \frac{TRH}{TRH + D_H} \tag{S68}$$

$$g_7 = \frac{\alpha_{S2} \beta_S}{\alpha_S \beta_{S2}} \tag{S69}$$

$$TSH_z = \frac{\alpha_{S2} \beta_S}{\alpha_S \beta_{S2}} TSH = g_7 TSH \tag{S70}$$

the steady-state equation of $TSH$ is

$$TSH = p_4 \frac{g_7 TSH + D_S}{g_7 TSH(1+S_S) + D_S} \frac{1}{1 + L_S G_R \frac{T_{3N}}{T_{3N}+D_R}}. \tag{S71}$$

With the expression (S67) and the definition of the constants

$$p_5 = g_7(1+S_S) \tag{S72}$$

$$p_6 = p_3 + p_2 L_S G_R \tag{S73}$$

$$p_7 = D_R K_{M2} \tag{S74}$$

$$p_8 = p_4 g_7 p_3 \tag{S75}$$

$$p_9 = p_4 g_7 D_R K_{M2} \tag{S76}$$

$$p_{10} = p_4 D_S p_3 \tag{S77}$$

$$p_{11} = p_4 D_S D_R K_{M2}, \tag{S78}$$

one gets the following equation

$$TSH^2(G_{MCT8}\frac{T_{4,th}}{K_{MCT8}+T_{4,th}}p_5p_6+p_5p_7)+TSH(G_{MCT8}\frac{T_{4,th}}{K_{MCT8}+T_{4,th}}(D_Sp_6-p_8)+D_Sp_7-p_9)$$
$$-G_{MCT8}\frac{T_{4,th}}{K_{MCT8}+T_{4,th}}p_{10}-p_{11}=0. \quad (S79)$$

By defining some final constants

$$q_1=p_5p_6 \quad (S80)$$

$$q_2=p_5p_7 \quad (S81)$$

$$q_3=D_Sp_6-p_8 \quad (S82)$$

$$q_4=D_Sp_7, \quad (S83)$$

one obtains the more compact steady-state equation (S16)

$$TSH^2(G_{MCT8}\frac{T_{4,th}}{K_{MCT8}+T_{4,th}}q_1+q_2)+TSH(G_{MCT8}\frac{T_{4,th}}{K_{MCT8}+T_{4,th}}q_3+q_4-p_9)$$
$$-G_{MCT8}\frac{T_{4,th}}{K_{MCT8}+T_{4,th}}p_{10}-p_{11}=0. \quad (S84)$$

#### S6 Numerical Parameter Values

| Symbol | Description | Value | Origin |
|---|---|---|---|
| $TBG$ | Concentration of thyroxine-binding globulin | 300 nmol/l | Neubert (1977)/ well known reference value |
| $TBPA$ | Concentration of Transthyretin | 4.5 $\mu$mol/l | Neubert (1977)/ well known reference value |
| $IBS$ | Concentration of intra-cellular $T_3$-binding substrate | 8 $\mu$mol/l | Estimated form $TBG$-concentration, corrected for intra-cellular $T_3$-accumulation (according to values from M. T. Hays (1988)) |
| $TRH$ | $TRH$-concentration in hypophyseal portal system | 6.9 nmol/s | Rondell et al. (1988) |
| $G_H$ | Secretory capacity of the pituitary | 817 mIU/s | Calculated according to D'angelo et al. (1976) and Okuno et al. (1979) |
| $D_H$ | Damping constant ($EC_{50}$) of $TRH$ at the pituitary | 47 nmol/s | Le Dafniet et al. (1994) |
| $\alpha_S$ | Dilution factor for peripheral $TSH$ | $0.4\,\mathrm{l}^{-1}$ | Reciprocal value of the volume of distribution of 2.5 l (Dietrich, 2001) |
| $\beta_S$ | Clearance exponent for peripheral $TSH$ | $2.3 \cdot 10^{-4}\,\mathrm{s}^{-1}$ | Calculated from plasma half-life of 50 min (Odell et al., 1967; Li et al., 1995) |
| $L_S$ | Brake constant of long feedback | 1.68 l/µmol | Calculated from clinical data of hyperthyroid patients (Dietrich, 2001) |
| $G_T$ | Secretory capacity of thyroid gland | 3.4 pmol/s | Li et al. (1995) |
| $D_T$ | Damping constant ($EC_{50}$) at the thyroid gland | 2.75 mIU/l | Dumont and Vassart (1995) |
| $\alpha_T$ | Dilution factor for $T_4$ | $0.1\,\mathrm{l}^{-1}$ | Reciprocal value of the volume of distribution (Greenspan, 1997) |
| $\beta_T$ | Clearance exponent for $T_4$ | $1.1 \cdot 10^{-6}\mathrm{s}^{-1}$ | Calculated from plasma half-life of 7 days (Greenspan, 1997; Grussendorf, 1988) |
| $K_{M1}$ | Dissociation constant of 5'-deiodinase type I | 500 nmol/l | Greenspan (1997) |
| $\alpha_{31}$ | Dilution factor for peripheral $T_3$ | $2.6 \cdot 10^{-2}\,\mathrm{l}^{-1}$ | Reciprocal value of volume of distribution (Greenspan, 1997) |
| $\beta_{31}$ | Clearance exponent for peripheral $T_3$ | $8 \cdot 10^{-6}\,\mathrm{s}^{-1}$ | Calculated from plasma half-life of 24 h (Dietrich, 2001) |
| $G_{D2}$ | Maximum activity of 5'deiodinase type II | 4.3 fmol/s | Calculated form pituitary $T_3$-concentration (Doorn et al., 1985) |

| Symbol | Description | Value | Origin |
|---|---|---|---|
| $K_{M2}$ | Dissociation constant of 5'-deiodinase type II | $1\,\text{nmol/l}$ | Visser et al. (1983) |
| $\alpha_{32}$ | Dilution factor for central $T_3$ | $1.3 \cdot 10^5\,\text{l}^{-1}$ | Calculated from volume of distribution 7.6 µl (Dietrich, 2001) |
| $\beta_{32}$ | Clearance exponent for central $T_3$ | $8.3 \cdot 10^{-4}\,\text{s}^{-1}$ | Calculated from intra-cellular half-life of 15 min (Oppenheimer et al., 1967; Benvenga and Robbins, 1998) |
| $\alpha_{S2}$ | Dilution factor for pituitary $TSH$ | $2.6 \cdot 10^5\,\text{l}^{-1}$ | Calculated from volume of distribution (Dietrich, 2001) of $3.8\mu l$ |
| $\beta_{S2}$ | Clearance exponent for pituitary $TSH$ | $140\,\text{s}^{-1}$ | Estimated, corresponding to half-life of 5 ms (Dietrich, 2001) |
| $D_R$ | Damping constant for central $T_3$ | $100\,\text{pmol/l}$ | Lazar et al. (1990) |
| $G_R$ | Maximum gain of $TR\beta$-receptors | $1\,\text{mol/s}$ | Value unknown, normalized to 1 (magnitude of feedback is determined by $L_S$) (Dietrich, 2001) |
| $S_S$ | Brake constant of ultrashort feedback | $100\,\text{l/mIU}$ | Determined according to values from Kakita et al. (1984) |
| $D_S$ | Damping constant for $TSH$ inside the pituitary | $50\,\text{mIU/l}$ | Determined according to values from Kakita et al. (1984) |
| $K_{30}$ | Dissociation constant $T_3$-$TBG$ | $2 \cdot 10^9\text{l/mol}$ | Li et al. (1995) |
| $K_{31}$ | Dissociation constant $T_3$-$IBS$ | $2 \cdot 10^9$ l/mol | Value unknown, adapted to extra-cellular dissociation constant (Dietrich, 2001) |
| $K_{41}$ | Dissociation constant $T_4$-$TBG$ | $2 \cdot 10^{10}$ l/mol | Li et al. (1995) |
| $K_{42}$ | Dissociation constant $T_4$-$TBPA$ | $2 \cdot 10^8\text{l/mol}$ | Li et al. (1995) |
| $\tau_{0S}$ | Peripheral delay for $TSH$ | 120 s | Derived from circulation time Dietrich (2001) |
| $\tau_{0S2}$ | Pituitary delay for $TSH$ | 3240 s | Derived from period of $TSH$-pulses (data from Greenspan et al. (1986)) |
| $\tau_{0T}$ | Delay for $T_4$ | 300 s | Estimated according to circulation and diffusion times (Dietrich, 2001) |
| $\tau_{03Z}$ | Delay for pituitary $T_3$ | 3600 s | Derived from Larsen (1982) |
| $\alpha_{th}$ | Dilution factor for $T_{4,th}$ | $250\,\text{l}^{-1}$ | Based on an assumed volume of distribution of 4 ml |
| $\beta_{th}$ | Clearance Exponent for $T_{4,th}$ | $4.4 \cdot 10^{-6}\text{s}^{-1}$ | Calculated from plasma half-life of 44 h (Di Cosmo et al., 2010) |
| $k_{Dio}$ | Stimulation constant of thyroidal D1 and D2 | 1 mIU/l | Berberich et al. (2018) |
| $K_{MCT8}$ | Michaelis-Menten constant MCT8 | $4.7 \cdot 10^{-6}$ mol/l | Friesema et al. (2003) |
| $G_{D1}$ | Maximum activity of 5'deiodinase type I | Compare main part | Fitted to real measurements |
| $G_{T3}$ | Maximum activity of direct $T_3$ synthesis | Compare main part | Fitted to real measurements |
| $G_{MCT8}$ | Maximum activity of MCT8 | Compare main part | Fitted to real measurements |
| $k_l$ | Linear approximation constant membrane transporters | Compare main part | Fitted to real measurements |
